# Impact of clade specific mutations on structural fidelity of SARS-CoV-2 proteins

**DOI:** 10.1101/2020.10.20.347021

**Authors:** Souradip Basu, Suparba Mukhopadhyay, Rajdeep Das, Sarmishta Mukhopadhyay, Pankaj Kumar Singh, Sayak Ganguli

## Abstract

The SARS-CoV-2 is a positive stranded RNA virus with a genome size of ~29.9 kilobase pairs which spans 29 open reading frames. Studies have revealed that the genome encodes about 16 non-structural proteins (nsp), four structural proteins, and six or seven accessory proteins. Based on prevalent knowledge on SARS-CoV and other coronaviruses, functions have been assigned for majority of the proteins. While, researchers across the globe are engrossed in identifying a potential pharmacological intervention to control the viral outbreak, none of the work has come up with new antiviral drugs or vaccines yet. One possible approach that has shown some positive results is by treating infected patients with the plasma collected from convalescent COVID-19 patients. Several vaccines around the world have entered their final trial phase in humans and we expect that these will in time be available for application to worldwide population to combat the disease. In this work we analyse the effect of prevalent mutations in the major pathogenesis related proteins of SARS-COV2 and attempt to pinpoint the effects of those mutations on the structural stability of the proteins. Our observations and analysis direct us to identify that all the major mutations have a negative impact in context of stability of the viral proteins under study and the mutant proteins suffer both structural and functional alterations as a result of the mutations. Our binary scoring scheme identifies L84S mutation in ORF8 as the most disruptive of the mutations under study. We believe that, the virus is under the influence of an evolutionary phenomenon similar to Muller’s ratchet where the continuous accumulation of these mutations is making the virus less virulent which may also explain the reduction in fatality rates worldwide.

## Introduction

What started off as an unusual outbreak of pneumonia in Wuhan China has reached the farthest corners of the world as a global pandemic. Till date there are over 60 million people affected by this novel strain of severe acute respiratory syndrome (SARS) virus and has been named as SARS-CoV-2 with the disease named as COVID-19. Genomic structures and phylogenomic studies reveal that the causal agent of COVID-19 belongs to genera Beta coronavirus. It is similar to the human Beta coronaviruses (SARS-CoV-2, SARS-CoV and MERS-CoV) but there are important differences at the genotypic and phenotypic levels which ultimately influence their pathogenesis. The SARS-CoV-2 contains a single stranded positive sense RNA encapsulated in a nucleoprotein matrix. The RNA genome of SARS-CoV-2 consists of approximately 29,800 nucleotides, encoding for 29 known proteins, though we find the expression of 28 to be constant with one remaining unexpressed. Each of the 28 proteins that are reported to be encoded perform different functions in the viral pathogenesis cascade though we are still not fully aware of the plethora of functions that they perform (Zeng et al., 2004). Four proteins make up the viral structure of which the most studied and important protein is the S protein or spike protein since they are the regulators of viral entry into the host. This protein binds to the angiotensin converting enzyme 2 (ACE2) receptor and gains entry into the host cell. A pair wise amino acid comparison using the amino acid sequences reveal that the spike proteins of SARS-CoV-2 share more than 80% sequence homology to SARS-CoV. Conservation is found in residues which interact with the ACE2 receptor; however, the other residues are a bit different with a few insertions as well. Till date the reports indicate that the interaction of the spike protein of SARS CoV-2 is much stronger as compared to the previous coronaviruses (Liu et al., 2004). A non-peer-reviewed preprint posted on medRxiv hypothesises that since ACE2 is overexpressed in the lung tissue of people with chronic respiratory disorders, they are likely to be more affected by this prevalent strain of the virus (Pinto et al., 2020). However, the infectivity of the virus is not restricted to its affinity for ACE2. If we dig deeper into the structure of the S protein we shall find that they are composed of two segments, S1 and S2 that require a double cleavage to expose the peptide which initiates the fusion with ACE2. The first cleavage is mediated by a host cell encoded enzyme furin. This ability to get human enzymes to do its work increases its infective ability as well. Few non-structural proteins of the virus are involved in the evasion of the host immune system and also function towards the assembly and replication of the virus. These proteins are expressed as part of two large polyproteins which are produced from a ribosomal frameshift event and are cleaved by the protease into 16 different proteins each with specific functions (Seah et al., 2020). The rest of the proteins are the accessory proteins which are technically non-essential as they are not required for viral replication in vitro and are the least well understood of the total proteome (Kim et al., 2020).

There have been several reports regarding the variation in severity and lethality of Coronaviruses. NL63, 229E, OC43, and HKU1 cause mild respiratory problems in humans and three others (MERS-CoV, SARS-CoV including the newly emerged SARS-CoV-2) have been observed to inflict severe respiratory syndromes (Veriti et al., 2020; WHO 2020; WHO 2020 [MERS-CoV]). SARS-CoV-2 infection was first reported from Wuhan, China, on 24th December 2019 (Veriti et al., 2020) and in less than three months, on 11th March 2020, WHO declared COVID-19 as a pandemic (Li, 2016). By May, 2020, 213 countries had been already affected with almost 2.5 lakhs fatal cases (He et al., 2004).Thus it was clearly visible that the infectivity of SARS-CoV-2 was much higher than SARS-CoV or MERS-CoV (Folis et al., 2006), though the fatality rate (0.9-3.3%) is still substantially lower than that of SARS-CoV (11%) and MERS-CoV (34%) (Tang et al., 2020).

Since this is a new pathogen, researchers worldwide are looking towards both primary and secondary prevention strategies in the form of vaccination as well as an over the counter medication which can be afforded by citizens around the world. Whatever strategy is being envisaged, the threat lies in the appearance of new strains with resistant mutations to vaccines and therapeutic agents. As a large proportion of humanity is already infected and we have no strategy in place currently to identify the asymptomatic carriers due to the lack of infrastructural and manpower resources around the world, there is high possibility of the successful escape mutations in the genome of the virus. Very few studies till date have focused on mapping the mutations on the important protein components of SARS-CoV-2. During the preparation of this manuscript we came across two studies which attempt to map the effects of significant mutations on the protein structure and stability. Benvenuto et al (2020) reports in more recent isolates of SARS CoV-2,the presence of two mutations affecting the non-Structural Protein 6 (NSP6) and the Open Reading Frame10 (ORF 10) in adjacent regions. Amino acidic change stability analysis suggests both mutations could confer lower stability of the protein structures. Namely, at amino acid position 3691 (corresponding to NSP6 position 37), most of the SARS-CoV-2 sequences have a leucine residue while some more recent sequences from Asian, American, Oceanian and European isolates show phenylalanine. At the amino acidic position 9659 (corresponding to ORF 10 position 3 to 4), most of the SARS-CoV-2 sequences have an arginine residue while some sequences from Australian and American isolates have a histidine residue. In the second study by Bhattacharya et al (2020) the 614 amino acid position in the S protein was found to be conserved among species (Bat, Pangolin, Civet and Human SARS-CoV and Human SARS-CoV-2), until the A2a subtype defining mutation occurred (D614G) (Bhattacharya et al., 2020). Authors report that an additional serine protease (elastase) cleavage site around S1-S2 junction was introduced due to D614G mutation in SARS-CoV-2. The Glycine at 614 (aa) is predicted to be the nearest substrate site for elastase to perform proteolytic cleavage in the adjacent residue.

This study focuses on mapping the effects of the clade specific mutations identified in the orf8, nsp2, nsp4, nsp6, nucleocapsid protein (N) and Spike protein (S) on the conformation and stability of the structures. It also predicts the possible drug binding sites, along with the druggability of these protein structures and the conformational epitopes of B cell, T cell, MHC -I and MHC –II alleles.

## Materials and Method

### IDENTIFICATION OF THE CLADE SPECIFIC MUTATIONS

Clade specific mutations were identified using literature mining from the various SARS-COV2 dedicated resources at PUBMED and PUBMED central and we identified a list of prevalent mutations across the major pathogenesis related proteins of the virus (Table 1). Though currently there are reports of other mutations that have been identified by large scale viral genomic sequencing studies undertaken globally, (Mercatelli and Giorgi 2020), the following dataset can be considered to be comprised of the initial founder mutations which resulted in the variations in the strain of the virus before the international borders were closed down (Callaway, 2020) following which the selection pressure on the virus became more country and population specific. Following the identification of the mutations we utilised the NCBI Genome Browser to extract the protein sequences using the Wuhan strain as the ancestral strain. Using a generic sliding window approach we were able to pinpoint the sites of mutations and we created two data sets - wild type protein sequences which were directly the ones from the Wuhan strain and the Mutant Data Set - which consisted of the protein sequences carrying the mutations in them

**Table 1:** List of clade specific mutations in the major pathogenesis proteins of SARS-CoV.

| **Mutation** | **Clade** | **Protein** |
| --- | --- | --- |
| N - S202N | B4 | Nucleocapsid protein (N Protein) |
| ORF1a - V378I | A3 | nsp2 protein |
| ORF1a - A3220V | A7 | nsp4 protein |
| ORF1a - L3606F | A3 | nsp6 protein |
| ORF1a - L3606F | A1a | nsp6 protein |
| ORF3a - G251V | A1a | ORF3a protein |
| ORF8 - L84S | B | orf8 protein |
| ORF8 - L84S | B1 | orf8 protein |
| ORF8 - L84S | B2 | orf8 protein |
| ORF8 - L84S | B4 | orf8 protein |
| S - D614G | A2 | Surface Glycoprotein (S Protein) |
| S - D614G | A2a | Surface Glycoprotein (S Protein) |

### SEQUENCE BASED ANALYSIS OF PROTEINS

#### Prediction of physicochemical properties of the protein

To understand the exact conformational behaviour of a protein, physical and chemical parameters of the wild type and mutant protein sequences were analysed using the ProtParam protein analysis tool of Expasy web server (https://web.expasy.org/protparam/) and PEPstat protein sequence statistical tool of EMBL-EBI web server (https://www.ebi.ac.uk/Tools/seqstats/emboss_pepstats/),to determine molecular weight, theoretical pI, extinction coefficient (Gill & von Hippel, 1989), estimated half-life (Bachmair et al., 1986), instability index (Guruprasad et al., 1990), aliphatic index (Ikai, 1980), grand average of hydropathy (Kyte & Doolittle, 1982), and isoelectric point (Harrison, 2000).

#### Prediction of Secondary structure, Accessible Surface Area and Intrinsic Disorder of the protein

The secondary structure of the wild type and the mutant proteins along with their degree of disordered residues and accessible surface area was predicted using the primary sequence of the protein. These were submitted as input to the NetSurfP - 2.0 server (http://www.cbs.dtu.dk/services/Net-SurfP/) (Klausen et al., 2019), which utilizes a neural network based approach trained on established protein structures as test sets. The accuracy of prediction is estimated to be 85% when compared to estimated structures. Since the degree of disordered residues in a protein has its impact on overall structural and functional stability of a protein, the data obtained from NetSurfP was cross-checked using the IUPred 2a server (https://iupred2a.elte.hu/) (Mészáros et al., 2018).

#### Prediction of Epitope Binding sites

The possible epitopes were predicted for the 7 wild type and their corresponding mutant proteins selected for the study. Linear B cell epitopes were predicted by using ABCpred (crdd.osdd.net/Raghava/abcpred/) and LBtope (webs.iiitd.edu.in/Raghava/lbtope/protein.php) servers. ABCpred requires an amino acid sequence as input and generates epitopes using an Artificial Neural Network (ANN) (Janahi et al., 2017). A window length of 16 amino acids and threshold of 0.51 were considered for generating the results (Kumar et al., 2018). “LBtope: Linear B cell epitope prediction server” was also used to reveal B cell epitopes of viral origin with a threshold probability score of 60% and above. The T cell epitopes of viral proteins binding to class I and class II Major histocompatibility complex molecules were mined using NetCTL 1.2 server (http://www.cbs.dtu.dk/services/NetCTL/) and Propred (https://webs.iiitd.edu.in/raghava/propred/). Viral peptides capable of binding to cytotoxic T lymphocytes and MHC I molecules were recognized using NetCTL 1.2 server which facilitates the prediction by combining the scores for proteasomal cleavage, TAP transport efficiency and MHC I binding (Larsen et al., 2007). The single letter code of amino acid sequence was entered in the user interface, the C terminal cleavage, weight on TAP transport efficiency and threshold for epitope identification was set at values of 0.1, 0.05 and 0.75 respectively. The viral proteins were screened against 12 most recognized supertypes of HLA alleles. The results were generated with a specificity of 97% and sensitivity of 80%, by integrating the scores based on multiple parameters as weighted sum with a relative weight on MHC I binding. For the prediction of MHC class II epitopes, Propred uses quantitative matrices methods which scan the antigen sequence against a profile of HLA-DR binding pockets, each having a unique quantitative representation of the amino acid interactions within the peptide binding cleft (Sturniolo et al., 1999). Amino acid sequence was provided as input at threshold value of 3, which enabled us to view the top 3% of best scoring peptides with respect to their probability of binding to different Human Leukocyte Antigen (HLA) molecules. Epitopes were predicted for all the 51 HLA-DR alleles that account for 90% of the MHC molecules on antigen presenting cells (Palanisamy & Lennerstrand, 2017).

#### Prediction of Changes in Protein Stability

The impacts of the mutations on the stability of the proteins were estimated using a sequence based approach at the I-Mutant 2.0 (Capriotti et al., 2005) web server. This is a program which classifies the effects of a particular mutation using a Support vector machine based approach. The training set of the tool was the Protherm database (Bava et al., 2004). The temperature and pH threshold used for this analysis were 25 and 7 respectively.

#### Generation of 3D structures for the wild type and mutant proteins

No complete crystal structures of these proteins are currently available in the Protein Data bank. As a result, ab initio modeling was performed using the I-TASSER server (https://zhanglab.ccm-b.med.umich.edu/I-TASSER/)(Roy et al., 2010) and the resultant structures were subjected to molecular simulation for 40ns using a GROMOS96 43a2 force field in Gromacs (Bekker et al., 1993; Abraham et al., 2015). Following the simulation, the structures were analysed for torsional/conformational stability from the Ramachandran plots of the models using the MOLPROBITY server (Williams et al., 2018). The integrity and validity of the 3D structure of the generated homology models were verified by Q Mean analysis.

#### Generation of Mutant Structures from wild type homology models

The variants prioritised by the sequence based analysis were inserted within the wild type models by using the MUTATE tool of DeepViewv4.1.0 (Guex & Peitsch, 1997) following which they were simulated and validated using the protocols followed for the wild type proteins.

#### Analysis of Importance of Position of mutation from an evolutionary structural perspective

The degree of evolutionary conservation of an amino acid in a protein is a signature of the balance between its natural tendency to mutate and the overall need for it to be retained in its original form to maintain the structural integrity and function of the macromolecule. In our analysis we used the ConSurf web server (http://consurf.tau.ac.il) (Ashkenazy et al., 2016), for predicting the evolutionary pattern of the amino acids of the macromolecule to reveal regions that are important for structure and function. The structures were submitted as inputs from which the sequences were extracted by the server and top 50 homologues were identified using three PSI - BLAST iterations against the UNIPROT database. Redundant homologous sequences were removed using the CD-HIT clustering method (Li & Godzik, 2006; Fu et al., 2012). The resulting sequences were then aligned using MAFFT (Katoh & Standley 2013) and the generated multiple sequence alignment (MSA) was used to generate the phylogenetic tree. The tree and the MSA were then considered together and the Rate4Site algorithm (Pupko et al., 2002) was used to calculate the position-specific evolutionary rates using an empirical Bayesian methodology (Mayrose et al., 2004). The raw data obtained was then normalized and grouped into nine conservation grades where 1 represented the rapidly evolving positions, 5 denoted positions with intermediate rates, and 9 was the threshold for the most evolutionarily conserved position. These were then mapped on the 3D structure on a position specific basis. We performed this analysis using both the Wild type and mutant structures.

#### Prediction of Binding pockets and Druggability

The alterations in potential ligand-binding sites and druggable pockets, in the wild type and mutant proteins were analysed using DoGSiteScorer (https://proteins.plus/) web server. The following parameters were assessed:

- Drug score- The total drug score was obtained by summing up the drug scores for each of the predicted pockets in the wild type and mutant structures. Increase or decrease in the total drug score was an indication towards the alteration of the pocket conformation and druggability.
- The number of ligand-binding pockets - The number of ligand-binding pockets in the wild type structure was considered as the reference and gain or loss of pockets in the mutant structure was considered for comparison.

#### Analysis of Structural Variation

Post simulation, structures of wild type and mutant pairs of each protein were aligned using TmAlign tool (Zhang & Skolnick, 2005). TmAlign performs structural alignment by superimposing structures, it performs a pair wise alignment for individual residues of protein. The output is provided in terms of Root Mean Square Deviation (RMSD), TmAlign score and aligned residues among structures. Here RMSD value proves to very critical when it comes to structural variations, higher the RMSD value higher are the differences among structures under consideration. In case of this study, higher the RMSD between structures, more were implications of mutations in terms of structural distortion. The molecular structures were visualized using Chimera.

#### Ranking of Mutations using a binary scoring scheme

Each sequence based and structural parameter analysed as a part of this study was given equal weightage since, the dataset was time limiting. A simple binary coding scheme was devised which was as follows: Alteration in value of parameter from wild type to mutant = −1, when the parametric value was unchanged for wild type and mutants then the value was “0” indicative of a neutral impact of the mutation. Using this scheme, a matrix was prepared and each mutation was provided with a cumulative score. This cumulative score was then arranged in a descending order and the mutation with the lowest score (most negative score) was considered to be the most disruptive.

## Results

### Determination of Protein Sequences from Genome

Using the methods outlined above we identified the wild type and mutant sequences which would be used subsequently (Table 2). Throughout the course of results and discussion we would be using the names of the proteins for easier understanding.

**Table 2:** Links and accession numbers of wild type protein sequences used in the study.

| PROTEIN | ACCESSION NUMBERS | LINK |
| --- | --- | --- |
| Nucleocapsid protein (N Protein) | >YP_009724397.2<br>nucleocapsid phosphoprotein | https://www.ncbi.nlm.nih.gov/nuccore/1798174254 [Link to Ancestral Clade] |
| nsp2 protein | >YP_009725298.1 |  |
| nsp4 protein | >YP_009725300.1 |  |
| nsp6 protein | >YP_009725302.1 |  |
| ORF3a protein | YP_009724391.1 |  |
| orf8 protein | YP_009724396.1 |  |
| Surface Glycoprotein (S Protein) | YP_009724390.1 |  |

### SEQUENCE BASED ANALYSIS

#### Physico- chemical properties of the protein

Physicochemical parameters were analysed using the servers mentioned in the materials and method segment using default algorithm parameters [Supplementary Table 1]. The changes observed in each of the physico-chemical parameters for the set of seven proteins are summarized in the following table (Table 3)

**Table 3:**
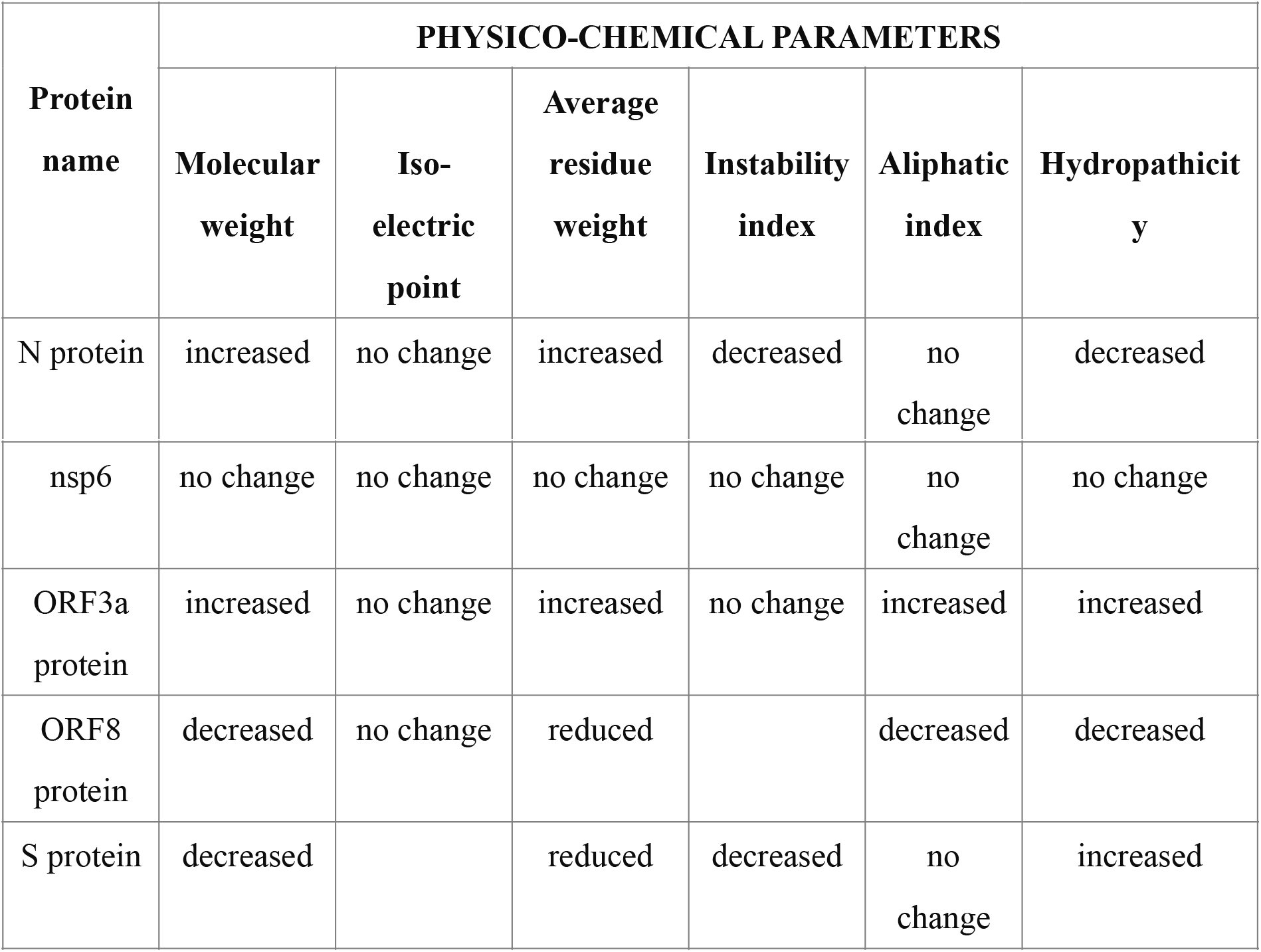
Alterations in physico-chemical parameters of proteins, post mutation.

#### Secondary structure, Accessible Surface Area and Intrinsic Disorder of the protein

Similar to the physico-chemical properties, no change was observed in accessible surface area and intrinsic disorder for nsp 6 protein from wildtype to mutant. (Figure 1a and 1b). However, distinct changes were observed in the accessible surface area and intrinsic disorder at the point of mutation for the following set of proteins - increase in accessible surface area in N protein (24.75 units), nsp 4 (25.72 units), ORF3a (23.46 units) and decrease in ORF8 (18.21 units) and S protein (30.44 units)were noted. In case of ORF3a, distinct decrease of intrinsic disorder was observed to be 0.106 units from wildtype to mutant protein, at the point of mutation. Positions not involving the mutation were observed to have secondary structural changes and changes in the accessible surface area **[Supplementary Table 2]** Secondary structural changes were noted for all the mutations and protein under study except nsp6. Alterations mostly involved abolition of specific structures such as helices and beta sheets **(Table 4** and Figure 2 a, b, c and d).

**Table 4:** Descriptions of alterations in Secondary structures. [Beta sheet alteration refers to the either breaking of one large beta sheets into various small ones or conglomeration of various small ones to form a larger beta sheet.]

| SL NO. | PROTEIN | POSITION | CHANGE FROM WILDTYPE TO MUTANT PROTEIN |
| --- | --- | --- | --- |
| 1 | N Protein | 250 - 255 | Helix discontinued |
| 2 | nsp2 | 420 - 426 | Helix discontinued |
| 3 | nsp4 | 40 | Beta sheet alteration |
| 5 | ORF3a | 70 - 80 | Coil structure is converted to alpha helix |
| 6 | ORF3a | 180 - 185 | Disappearance of alpha helix |
| 7 | ORF3a | 190 | Shortening of beta sheet |
| 8 | ORF3a | 256 | Addition of a beta sheet |
| 10 | S Protein | 560 | Loss of beta sheet |
| 11 | S Protein | 675 - 690 | Beta sheet alteration |
| 12 | S Protein | 1205 - 1215 | Change in the pattern of helix |

### EPITOPE PREDICTION RESULTS

Epitopes harboured in the viral proteins, that elicit immune responses in the host, can serve as potent vaccine candidates in near future. Linear B cell epitopes, were ascertained using servers, ABCpred and LBtope. LBtope organizes the antigenic determinants in descending order of their probability scores. Epitopes with greater than 60% prediction scores were additionally screened for epitopes encircling the alteration site (Supplementary Table 3). The number of such epitopes and their mean prediction scores were evaluated in both wild type and mutant proteins. While the numbers of epitopes were identical in both normal and mutant forms for each of the seven proteins, the mean value of the prediction scores, showed variation in each case, suggestive of the mutation induced changes in binding efficiencies (TABLE 5 and Figure 3).

**Table 5:**
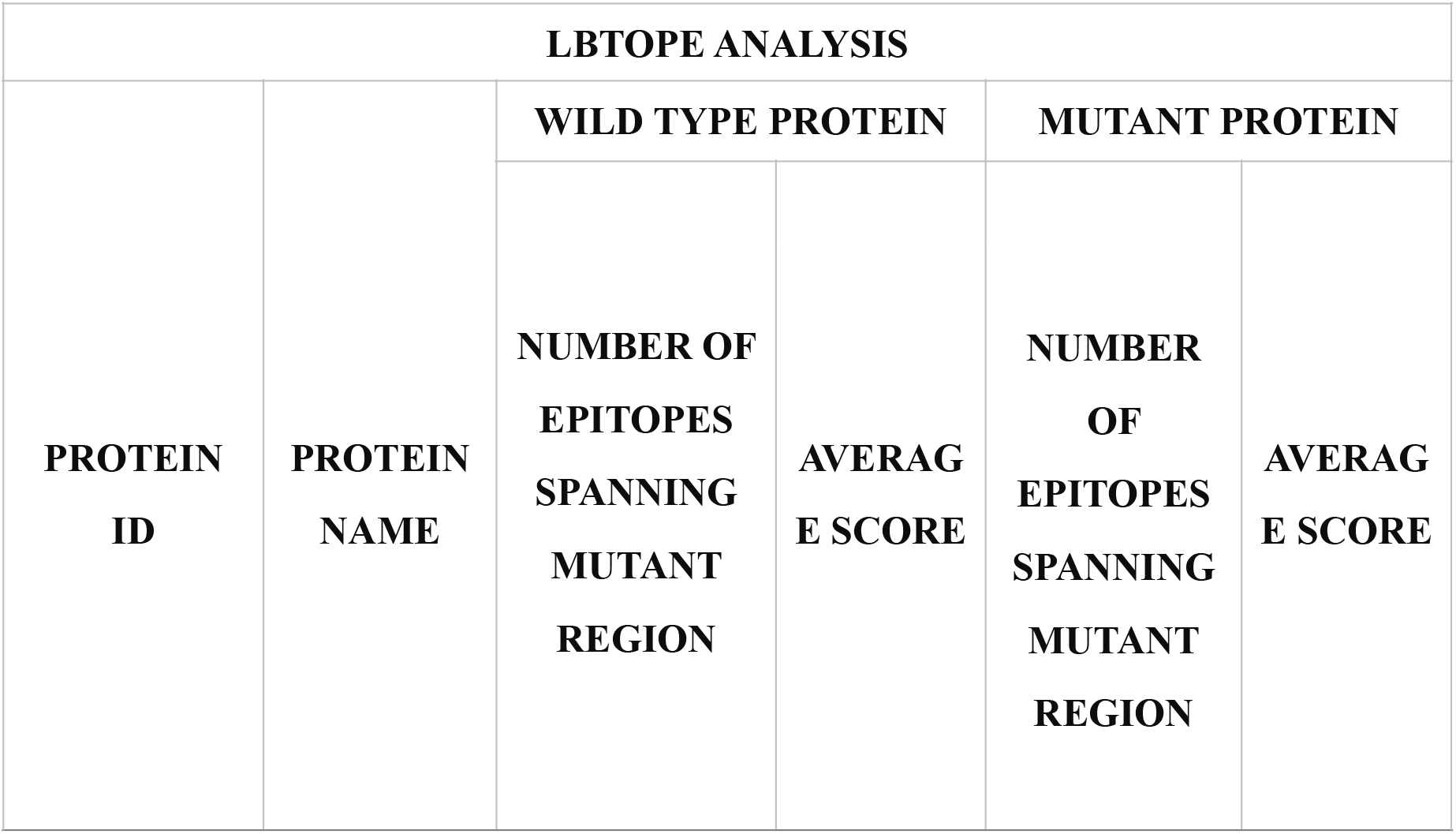
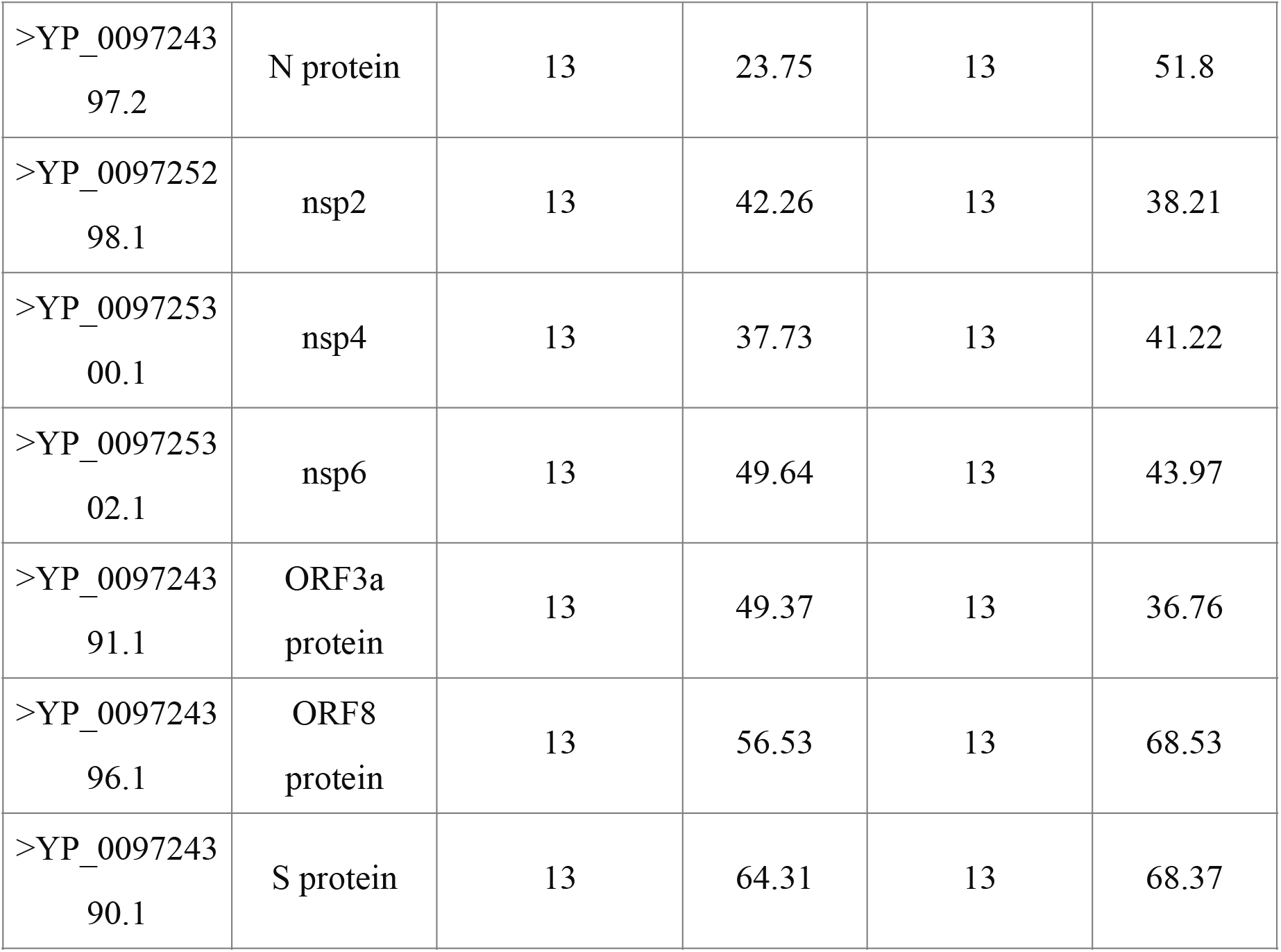
Variation in average score of epitopes for the set of seven proteins using LBtope.

For ABCpred, the antigenic determinants are ranked with respect to their binding probabilities, from highest to lowest, thus endorsing the best binder the first rank and so on. In this work, we have analysed the epitopes having any rank within 10 for each protein, enumerated their numbers and computed their average score for wild type and mutant forms (TABLE 6). We observed a change in the number of epitopes and their mean binding score for N protein and nsp4 proteins. While the number of epitopes remained unaltered, the mean score showed discrepancy for the ORF8 protein. An epitope holding the alteration site in wild type ORF8 protein had a binding score of 0.64, which was altered to 0.67 post mutation. Again, in proteins like nsp4 and N protein, mutation resulted in recognition of new epitopes covering mutation region, which was not predicted previously in wild type proteins. No fluctuation in epitope count and score was noted in case of ORF3a, S protein, nsp2 and nsp6 proteins (Figure 4).

**Table 6:** Comparison of linear B-cell epitopes between wt and mutant proteins using ABCpred.

| PROTEIN ID | PROTEIN NAME | WILD TYPE PROTEIN |  | MUTANT PROTEIN |  |
| --- | --- | --- | --- | --- | --- |
|  |  | NUMBER OF EPITOPES WITHIN TOP 10 RANK | MEAN PROBABILITY SCORE | NUMBER OF EPITOPES WITHIN TOP 10 RANK | MEAN PROBABILITY SCORE |
| >YP_00972439<br>7.2 | N protein | 21 | 0.869 | 22 | 0.868 |
| >YP_00972529<br>8.1 | nsp2 | 17 | 0.882 | 17 | 0.882 |
| >YP_00972530<br>0.1 | nsp4 | 21 | 0.869 | 22 | 0.866 |
| >YP_00972530<br>2.1 | nsp6 | 14 | 0.794 | 14 | 0.794 |
| >YP_00972439<br>1.1 | ORF3a protein | 12 | 0.884 | 12 | 0.884 |
| >YP_00972439<br>6.1 | ORF8 protein | 10 | 0.782 | 10 | 0.785 |
| >YP_00972439<br>0.1 | S protein | 28 | 0.891 | 28 | 0.891 |

NetCTL server was used at IEDB analysis resource for predicting the MHC class I T cell epitopes. Primary protein sequence was selected as input by us. Integration of proteasomal C-terminal cleavage and TAP transport efficiency were considered with default values, that is 0.15 and 0.05 respectively. We screened all the 12 supertypes for the study. During the analysis, A3 supertype of N protein; A26, B7, B3 supertypes of nsp2; A24, A26, B8, B27, B62 of nsp4; A1, A2, A24, A26, B27, B58, B62 supertypes of nsp6; A26, B44 supertypes of orf3a; A2, A24, A26, B62 supertypes of orf8; A1, A2, A3, B39, B58, B62 supertypes of S protein; showed variation in both wild type and mutant on the basis of average combined score **(Supplementary Table 4)**. Finally we summed up all the combined scores of all the epitopes of the individual proteins lying above their respective average score. The data of table Y was represented through a histogram (Figure 5).

Propred was used to predict MHC class II binding T cell epitopes from the set of 14 proteins. The top 3% of the best scoring epitopes generated from each protein were further shortlisted on the basis of the average score of the epitopes encompassing the mutation region in the wild type protein. The epitopes with probability scores equal to or greater than the average score was screened from the wild type and mutant proteins. The filtered epitopes were then, quantitatively and qualitatively compared and scrutinized between normal and mutant forms **(Supplementary Table 5)**. In case of ORF3a protein, 6 less epitopes were found to qualify the average score (38.91) in the mutant protein, however the number of epitopes spanning the mutation region remained unaltered. Contrasting results were obtained for ORF8 protein, where an epitope holding alteration segment recognized by the allele DRB1_0309 in wild type protein, remain unperceived by the same allele in the mutant protein. Also 4 less antigenic determinants were detected in the mutant protein above the standard mark. In case of the S protein, the mutation resulted in both abolishment and establishment of new epitopes enclosing the mutation site, recognized by several alleles in wild type and mutant proteins, pointing to the changes in binding capabilities emerging out of mutation. The nsp4 protein showed dissimilarities in the number of epitopes faring well above the cut off score, with 2 antigenic determinants (surrounding mutation site) remaining unrecognized by the same allele in the mutant protein and 4 epitopes (spanning mutation site) showing altered probability score. Changes in the number and binding specificity of epitopes and deviation of scores were also noted between wild type and mutant forms of nsp6 proteins. No changes in epitopes and their binding efficiencies were observed in case of the N protein and nsp2 protein (Table 7 and Figure 6).

**Table 7:** Quantitative estimation of MHC II binding T-cell epitopes in wt and mutant proteins, using Propred.

| PROPRED ANALYSIS |  |  |  |
| --- | --- | --- | --- |
|  |  | WILD TYPE PROTEIN | MUTANT PROTEIN |

| PROTEIN<br>ID | PROTEIN<br>NAME | NUMBER<br>OF<br>EPITOPES<br>COVERING<br>MUTANT<br>REGION | NUMBER<br>OF<br>EPITOPES<br>NOT<br>COVER-ING MU-TANT RE-GION | TOTAL<br>NUMBER<br>OF EPI-TOPES<br>ABOVE<br>AVERAG<br>E<br>SCORE | NUMBER<br>OF<br>EPITOPES<br>COVERIN<br>G<br>MUTANT<br>REGION | NUMBER<br>OF<br>EPITOPES<br>NOT<br>COVER-ING MU-TANT RE-GION | TOTAL<br>NUMBER<br>OF EPI-TOPES<br>ABOVE<br>AVERAG<br>E<br>SCORE |
| --- | --- | --- | --- | --- | --- | --- | --- |
| >YP_009724397.2 | N<br>protein | 0 | 25 | 25 | 0 | 25 | 25 |
| >YP_009725298.1 | nsp2 | 4 | 46 | 50 | 4 | 46 | 50 |
| >YP_009725300.1 | nsp4 | 6 | 40 | 46 | 4 | 44 | 48 |
| >YP_009725302.1 | nsp6 | 14 | 30 | 44 | 11 | 36 | 47 |
| >YP_009724391.1 | ORF3a<br>protein | 14 | 24 | 38 | 14 | 18 | 32 |
| >YP_009724396.1 | ORF8<br>protein | 1 | 49 | 50 | 0 | 46 | 46 |
| >YP_009724390.1 | S<br>protein | 27 | 11 | 38 | 29 | 6 | 35 |

**Table 8:** Impact of mutations on Epitope positioning and ASA values.

| Sl no. | Protein | Epitope positions which suffer secondary structural alterations (detailed in Table 4) | Change in accessible surface area |
| --- | --- | --- | --- |
| 1 | N Protein | 249,243 | PRESENT |
| 3 | nsp4 | 40 | PRESENT |
| 6 | ORF3a | 176, 184 | PRESENT |
| 7 | ORF3a | 190 | PRESENT |
| 8 | ORF3a | 243 | PRESENT |
| 12 | S Protein | 1206 | PRESENT |

#### Prediction of Protein Stability

Potential changes to stability of the proteins brought about by the mutations were analysed using the I-Mutant server. According to the algorithm of the server the range of DDG values for a Neutral mutation classification is : −0.5=<DDG=<0.5, while changes < †0.5 are classified as Large Decrease and changes > 0.5 are classified as Large Increase. Thus in our analysis we found that N protein, orf3a and S protein could be categorised as having large decrease with orf8a having the highest negative value of −2.87, while nsp 2 had large increase in the value. Alterations of nsp4 and nsp6 were predicted to be in the neutral change segment (Figure7).

#### Results of Structural Analysis

Stable structures were generated using ab initio modeling which were subsequently simulated [Figure 8a, b and c(i)]and their conformational accuracy was analyzed using Ramachandran plot and Q mean analysis. In the Ramachandran plot analyses **(Supplementary Figure 1)**it was revealed that all the mutations resulted in the accumulation of new outlier residues in the protein structure evaluation indicating that fresh regions of destability has been introduced into the structures. An interesting observation was that in case of nsp4 and orf 8a where though the numbers of outlier residues were exactly equal in case of wild type and mutant structures, 12 and 6 new outlier residues were incorporated as part of the total outlier number, for nsp4 and orf8a respectively. For orf 3a and nsp6 proteins there was a reduction in the total number of outliers from 30 in case of wild type to 22 in case of the mutant with 10 new outlier residues for orf3a while for nsp 6 the numbers were 21 and 13 respectively [Figure 8c(ii)].

QMeans values were found to differ in case of the wild type and mutant structures with significant changes observed in case of S protein, orf8a and orf 3a; while in case of nsp2 opposite trend was observed, though the differences in QMeans score was not very significant for nsp2 (Figure 9). All the fluctuation plots obtained following simulation [Figure 10 a, b c (i)] exhibited differences in the fluctuation profiles of mutant structures as compared to wild type indicative of randomness in conformation induced by the mutations. These were further reflected in their variation of average RMSF [Figure 10c(ii)].

To further confirm the variations induced as a result of the mutations we aligned the protein structurally and distinct differences were observed (Figure 11). Mutations led to differences in the protein structure of every mutant - wild type pair. Highest RMSD value was observed for S protein (5.67), while maximum number of mismatches was observed in case of N protein (24 out of 419). N Protein also had a high RMSD of 4.93. The results of structural alignments do suggest that mutations altered the protein structures substantially (Figure 12). The analysis of potential drug binding pockets and their druggability was further evaluated and all the proteins had alterations not only in the total number of drug binding pockets but also the average drug scores when wild type and mutant structures were compared. It was interesting to note that apart from nsp4 and orf8a all the other proteins, suffered a decrease in average druggability as a result of the mutations whereas the former two proteins exhibited an increase in potential druggability *(Figure 13 a and 13 b).* No significant variations were observed in our evolutionary analysis using CONSURF server.

#### Scoring of Mutations for Assessment of Impact

A simple binary scoring scheme was devised, where a parameter value alteration was indicated as −1 (indicating a change from wild type value) and unaltered parameter (indicating of no change from wild type parameter) was used for the analysis. For parameter values which involved decimal numbers, change was considered only if the first place after decimal had been altered **(Supplementary Table 6)**. Once the scoring scheme was populated with the corresponding values a heat map was generated to diagrammatically exhibit the contributions of each parameter under study towards the overall cumulative score obtained *(Figure 14 a).* The ranking of the mutations were then performed using a histogram in which the cumulative scores were plotted *(Figure 14b)*. From the histogram we were able to clearly predict the L84S mutation in ORF8a as the most disruptive of the mutations followed closely by nsp4 mutation (ORF1a 3220V) and the mutations in N protein (S202N); or f3A (G251V) and S protein (D614G). The lowest ranked mutations were those of nsp 6 (L3606F) and nsp2 (V378I).

## Discussion

### Physicochemical Analysis and Sequence Based Predictions

According to Kyte et. al. (1982) increased value of hydropathicity indicates the presence of more hydrophobic residues in the protein sequence and Lins et. al. (2003), points out that decrease in accessibility is more pronounced for hydrophobic than hydrophilic residues (Kyte &Doolittle, 1982; Lins et al., 2003). In our analysis S protein and ORF3a have increased hydropathicity and decreased accessible surface area which suggest that the mutation have resulted into increased hydrophobicity. Salahuddin and Khan (2009) have reported that increase in hydropathicity can be a direct determinant of increased pathogenicity for viral proteins. This observation is supported by the observations of (Sur et al., 2009). Both of these works involve analysis of mutations and genome variations with flu viral genomes. S protein acts as a cellular receptor to induce the fusion of viral and host membranes (Tan et al., 2004; Tan et al., 2005). It is important for the viral entry into the host cell and for inducing host immune responses. Sequence comparison of isolates from different infections show both viral S protein and ORF3a gets positively selected during evolution (Yin, 2020) implying that ORF3a plays an important role in the virus life-cycle and disease development. Also, according to Linding et. al. (2003) protein disorder is important for understanding protein function and protein folding pathway (Linding et al., 2003). In our study, ORF3a protein exhibits decreased disorderedness in its mutant form which indicates towards a change in the structural conformation and rigidity. In our study, maximum change in accessible surface area was observed in S protein whereas in case of ORF3a the intrinsic disorder was observed to decrease significantly which may affect the role of S protein’s functional interaction along with ORF3a. Ikai et. al. (1980), defined aliphatic index as the relative volume of a protein occupied by aliphatic side chains (alanine, valine, isoleucine, and leucine) of amino acids which show a correlation with increase in thermostability of proteins (Ikai, 1980). In our analysis nsp2, nsp4 and ORF3a were observed to show an increase in aliphatic index value from wild type to mutant which suggests probable increase in the thermostability of these proteins due to mutation. Interestingly ORF3a protein has a significant role in the functioning of S protein in the host system to increase infection which may be affected by the increased thermostability of ORF3a. Momen-Roknabadi et. al. (2008), in their study predicted that accessible surface area can be used to improve the prediction of secondary structure of a protein (Momen-Roknabadi et al., 2008). According to Lins et al (2003), different secondary structures correspond to different accessibility of residues. In their study, they found that random coils, beta sheets were more accessible folds with average 30% accessibility, while the alpha helices have 20% accessibility. In our analysis, we found a similar phenomenon where loss of beta sheet, conversion of random coils to alpha helices have led to decreased accessible surface area and loss of alpha helix have caused an increase in accessible surface area.

#### Epitope Prediction

Induction of adaptive immunity is largely dependent on recognition of viral epitopes by host B and T cell receptors. Site specific mutations in the mentioned set of seven proteins from Sars-COV2 were extensively analyzed for their effects on epitope recognition and binding efficiencies. While gain of function mutations can improve the binding affinities of epitopes to host lymphocyte receptors, loss of function mutations arising through positive selection can be crucial for B and T cell immune escape (Ramaiah et al., 2019). Each of the seven mutant proteins investigated, exhibited either change in free energies of binding or recognition by specific alleles, thus necessitating further exploration to evaluate their impact on immune escape and viral persistence. Besides, changes in the number of linear B-cell epitopes were spotted in proteins like nsp4 and N protein. In our study, we observed an epitope covering mutation region, recognized by A3 supertype in wild type of N protein but no changes in corresponding epitope was spotted by the same supertype in the mutant N protein (loss of function mutation); in contrast we spotted a mutation spanning epitope in mutant nsp4 protein perceived by A3 supertype, which was not detected by the same supertype in the wild type nsp4 protein (gain of function mutation). Another significant finding was the decline in the number of MHC II binding T-cell epitopes in the ORF8 and S protein, where the mutant protein was found to have 4 and 3 less immunogenic determinants respectively. Each of the seven proteins were assigned a score of either ‘-1’ or ‘0’, for each of the four computational tools used for epitope prediction, where ‘-1’ corresponds to any change in number or binding efficacy of antigenic determinants, that may have surfaced because of mutation and ‘0’ corresponds to no changes between wild type and mutant forms.

#### Structural Analysis

The RMSF value measures the deviation between the positions of particle i and some reference position. RMSD and RMSF values can be differentiated remarkably. RMSF is averaged over time, depicting a value for each particle i, while in case of RMSD, the average is taken over the particles, giving time specific values. Zhao et al (2015) have probed into the effect of mutations in the Zinc Binding B box1 domain protein (Zhao et al., 2015). In their study they have looked into the changes in the dynamics of the backbone atoms and also calculated root mean square fluctuation (RMSF) values at each time point of the trajectories of the native and mutant structures. They found that mutations induced higher RMSF which led them to suggest that the much larger side-chain of valine extends its steric hindrance to other zinc-binding residues. Supporting the increased flexibility, they also measured the RMSF values for second zinc-binding residues (C134, C137, H150, H159); which led to the final conclusion that: larger RMSF values indicate increased random motions of residues leading to disruption in the structural integrity of the proteins. In our analysis we have also observed a similar phenomenon with the increase in the RMSF values in mutant structures ultimately causing randomness in the 3D structures which is evidenced by their increased RMSD and more negative Q mean scores. The significance of Q mean scores have been extensively reported by de Carvalho and De Mesquita (2013) in their work on human superoxide disputes 2 where they compared wild type and mutant structures and reported the effect of mutations (de Carvalho & de Mesquita 2013). Using the TM align and Q mean methodology Datta et al (2015) have explored the effect of 124 single nucleotide polymorphisms of the ribonuclease L gene (RNASEL) in prostate cancer (Datta et al., 2015). All the SNP’s were mapped on the protein structure and TM Align was used to classify the most disrupting mutants. In our analyses we have been able to clearly show that mutations caused significant changes in protein structures where highest RMSD value was observed for S protein (5.67), while maximum number of mismatches were observed in case of N protein (24 out of 419). N Protein also had a high RMSD of 4.93. Thus from the structural comparisons of wild type and mutant proteins we were able to understand that almost all the mutations destabilised their native conformation.

#### Scoring and Final Prediction of Impact of Mutations

Mutations in all the proteins were first matched with the alteration data and it was found that direct effect of the mutation residue was associated with the changes in the Epitope site predictions (Table 6). They were also associated with alteration in the accessible surface area values as predicted thus implying that the mutations resulted in the changes in both structural and functional aspects of the protein structures.

Huskova et al 2017 have developed a scoring scheme for analysis of cancer driver mutations where they prioritized them using a simple scoring system. A score of 1 was added if the mutation was of the exposure-predominant type (likely introduced early in the assay) (Huskova et al., 2017). A score of 1 was added if the mutation was in a known human hotspot, if it was truncating or affected a splice site and so on. In our scoring scheme we prioritised the ability of the mutation to cause deviation in the structure of the mutant from the wild type and assigned scores as described in the materials and method segment. Our final scoring revealed that ORF8 protein mutation L84S was the most disruptive of the mutations under study. Incidentally L84S mutation was also the most prevalent of the mutations under study as it was present in four different clades B, B1, B2 and B4 clades. Incidentally Bhattacharya et al (2020) have reported the incidence of these subtypes in different countries (Bhattacharya et al., 2020). For example B subtype have been reported in large numbers from Australia(34), Belgium (10), China (63), Spain (66) UK (26) and USA (28); B1 has been reported from Australia (42), Canada (40), and USA (446); B2 has been identified from China (10) and Japan (4); while B4 finds its prevalence in Australia (5), China (5) and USA (27). These data suggest that this mutation was one of the earliest of the mutations that could have occurred naturally as it does not exhibit any population specificity and may be attributed to a random event. This data was however consistent with the predictions made in the I Mutant server which had assigned the most negative value for L84S and classified it as the most disruptive. This data justifies our scoring scheme and the parameters considered towards the prediction of the most disruptive mutation and thus can be explored further for future studies. Recent works suggest that the lowering of the stability of viral proteins as a result of accumulation of mutations (Banerjee et al., 2020) could be the reason behind the low National fatality rate in India. It may also be a phenomenon similar to Muller’s ratchet where this accumulation of mutations in fresh strains of the virus and lack of stabilising recombination events may result in the reduction in fitness of the virus as is being indicated by the effect of the mutations on protein stability.

## Supporting information

Ramachandran Plots Generated

Physicochemical Parameters

Total residue specifc values of ASA & DISORDER

LBtope prediction table

NetCTL prediction table

Propred Prediction Table

Scoring Scheme

## Acknowledgement

The authors acknowledge the contributions of Dr. Souvik Mukherjee, National Institute of Biomedical Genomics, Kalyani, West Bengal for his insightful discussions regarding the data acquisition techniques as well as overall comments during the preparation of the manuscript.

## List of Supplementary Material

1. Supplementary Table 1: Physicochemical parameter table

2. Supplementary Table 2: Total residue specific values of ASA and Disorder - [Single Table with multiple sheets for each individual protein]

3. Supplementary Table 3: LBtope prediction table

4. Supplementary Table 4: NetCTL prediction table

5. Supplementary Table 5: Propred predictionTable

6. Supplementary Table 6: Scoring Table for Impact Assessment

7. Supplementary Figure 1: Images of Ramachandran Analysis for Wild Type and Mutant Structures.

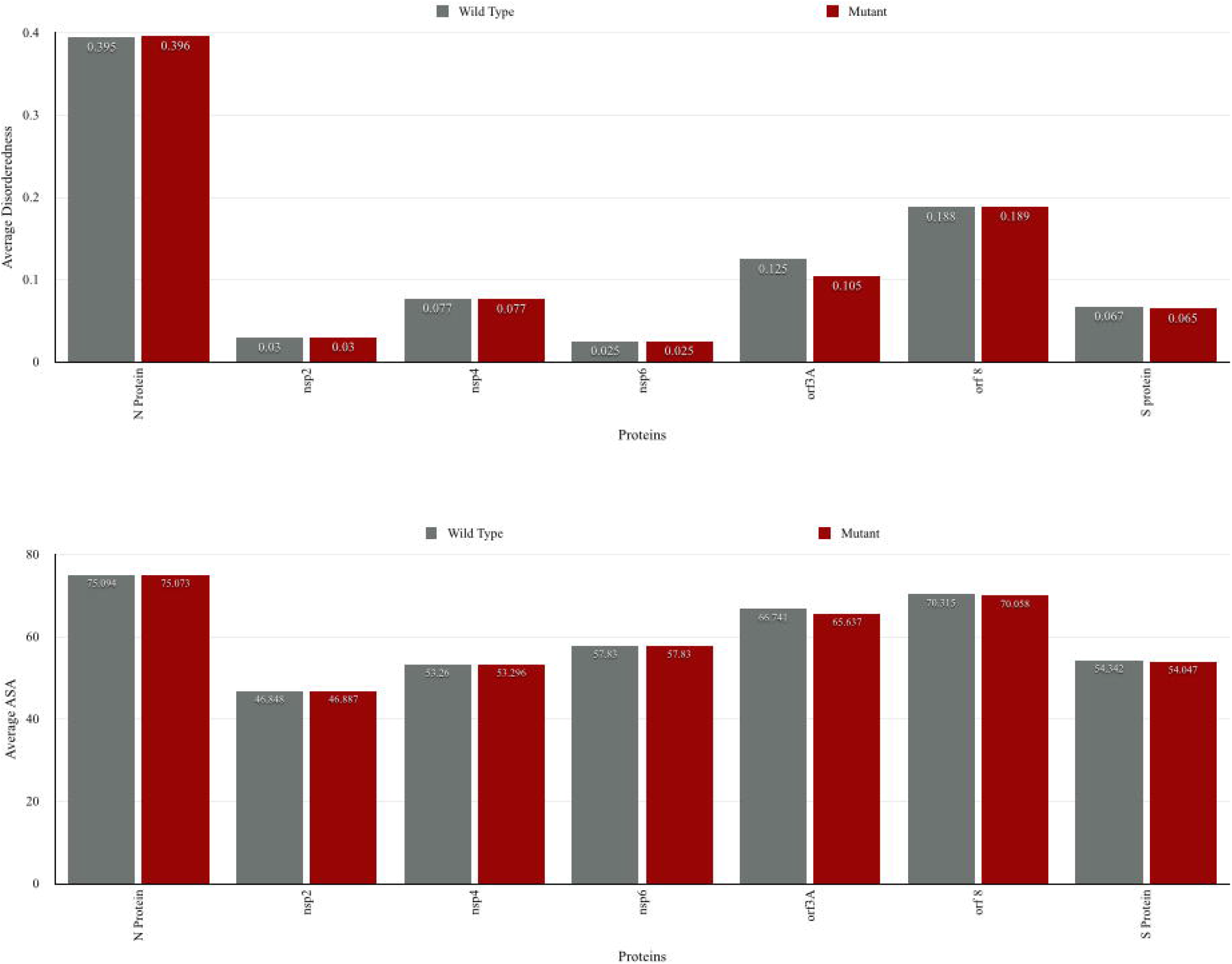
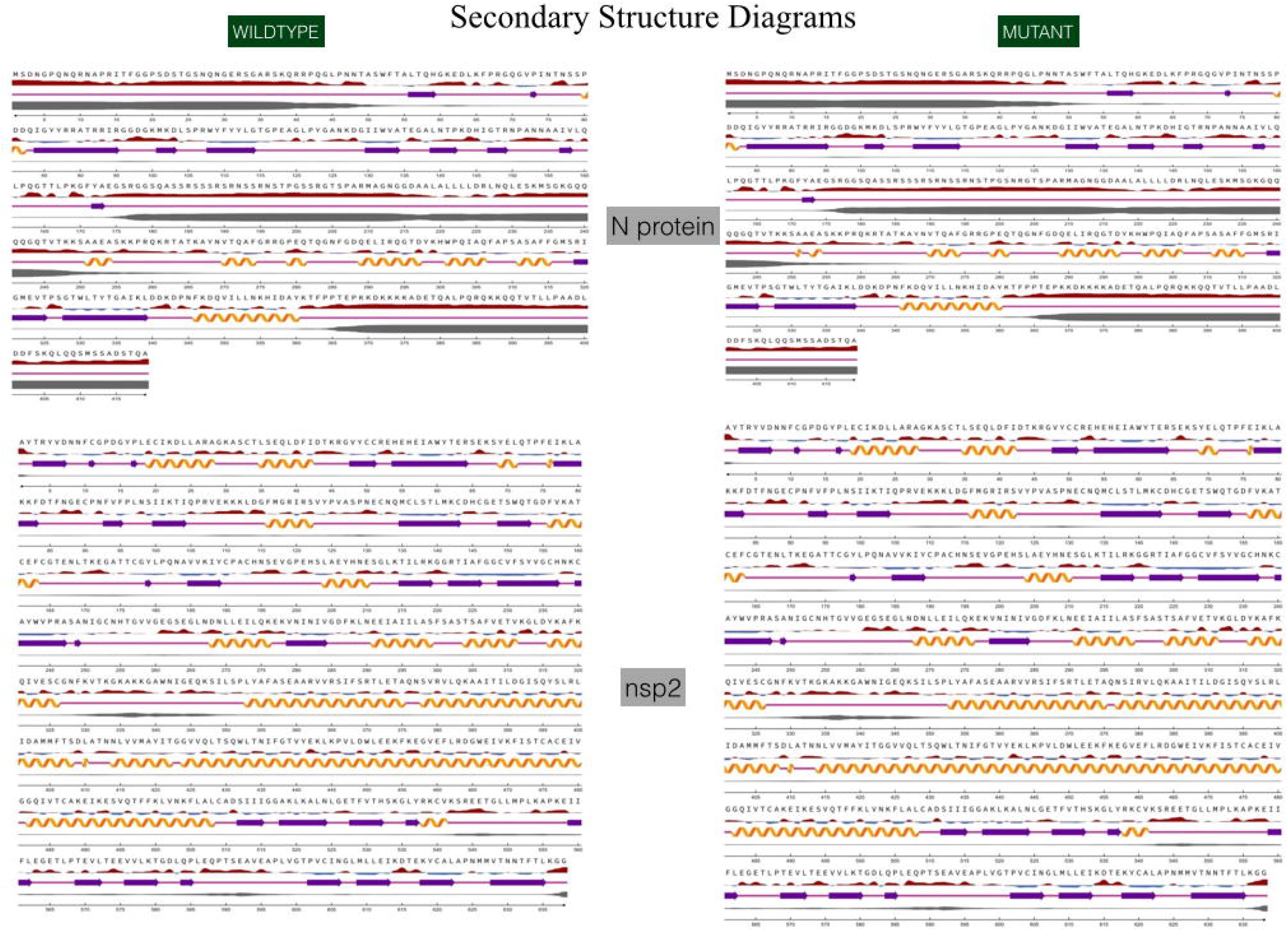
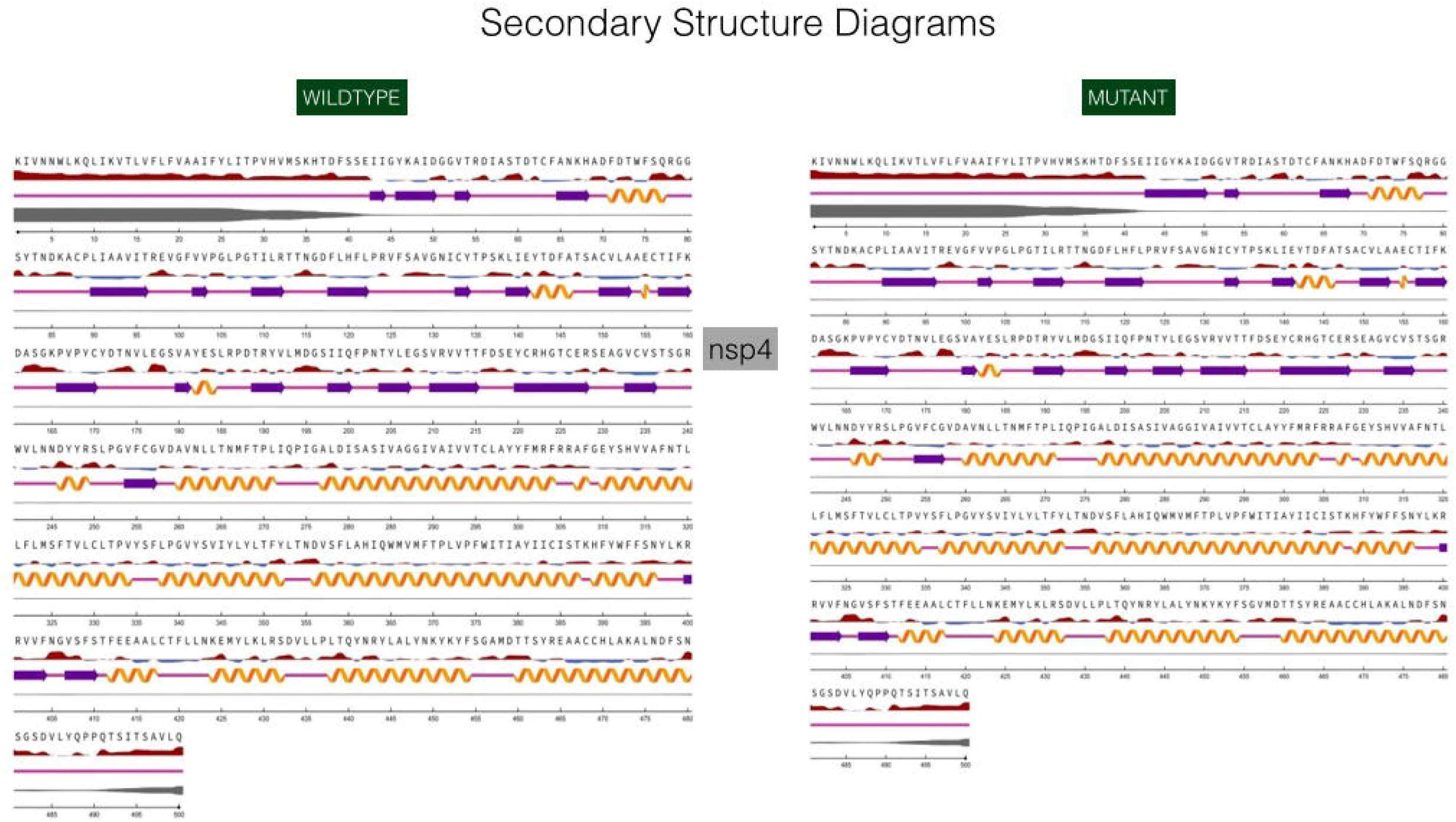
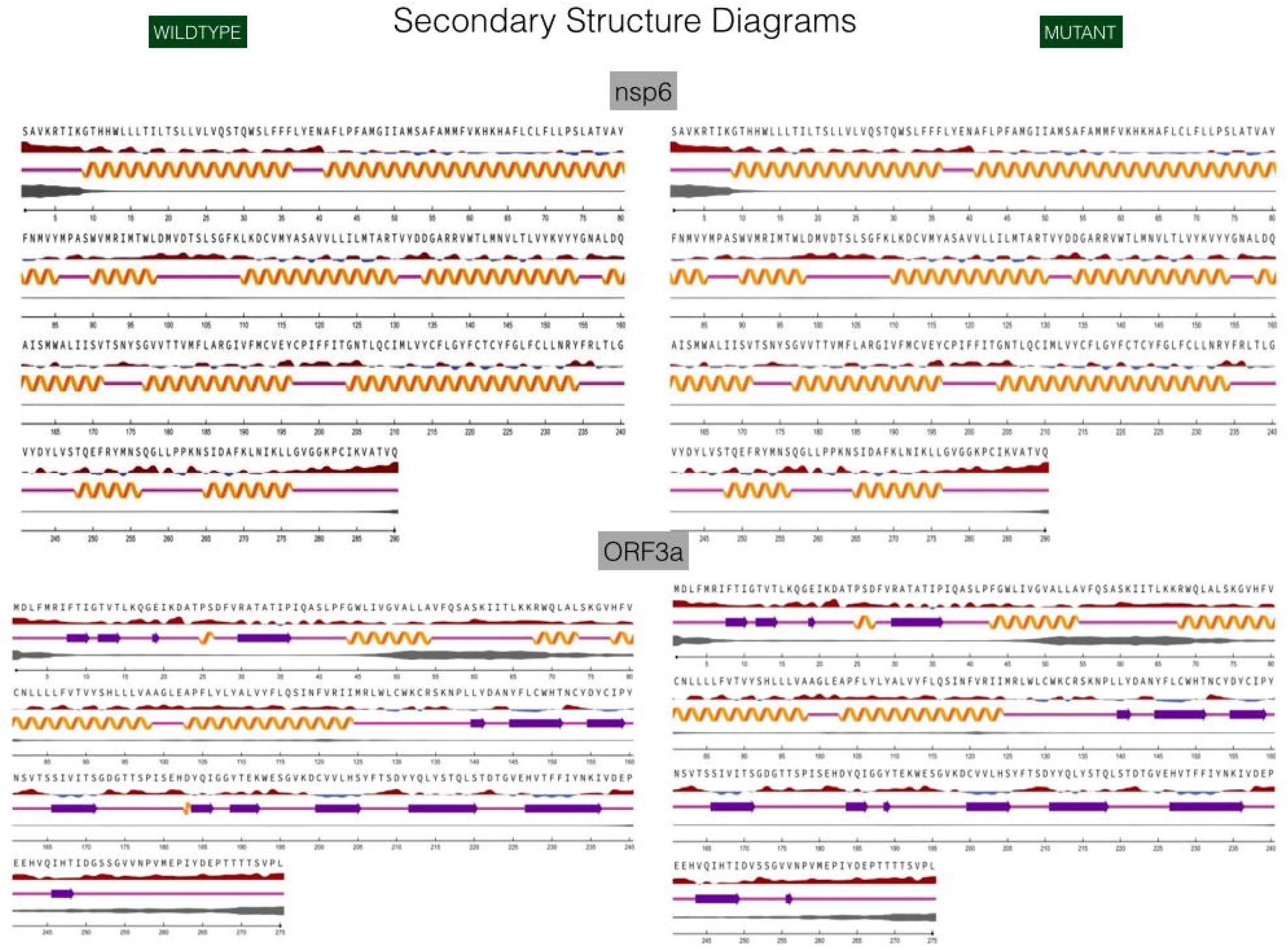
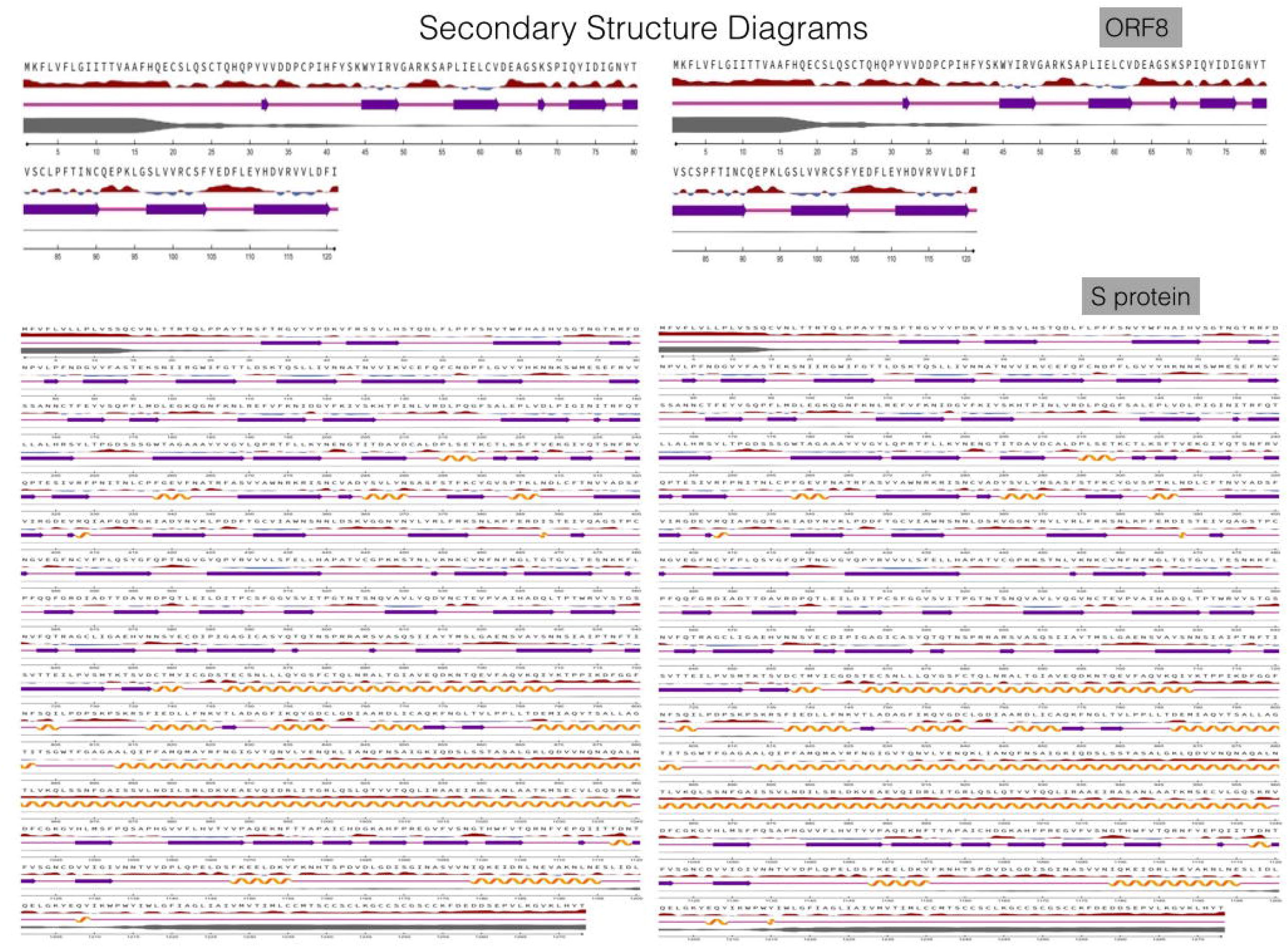
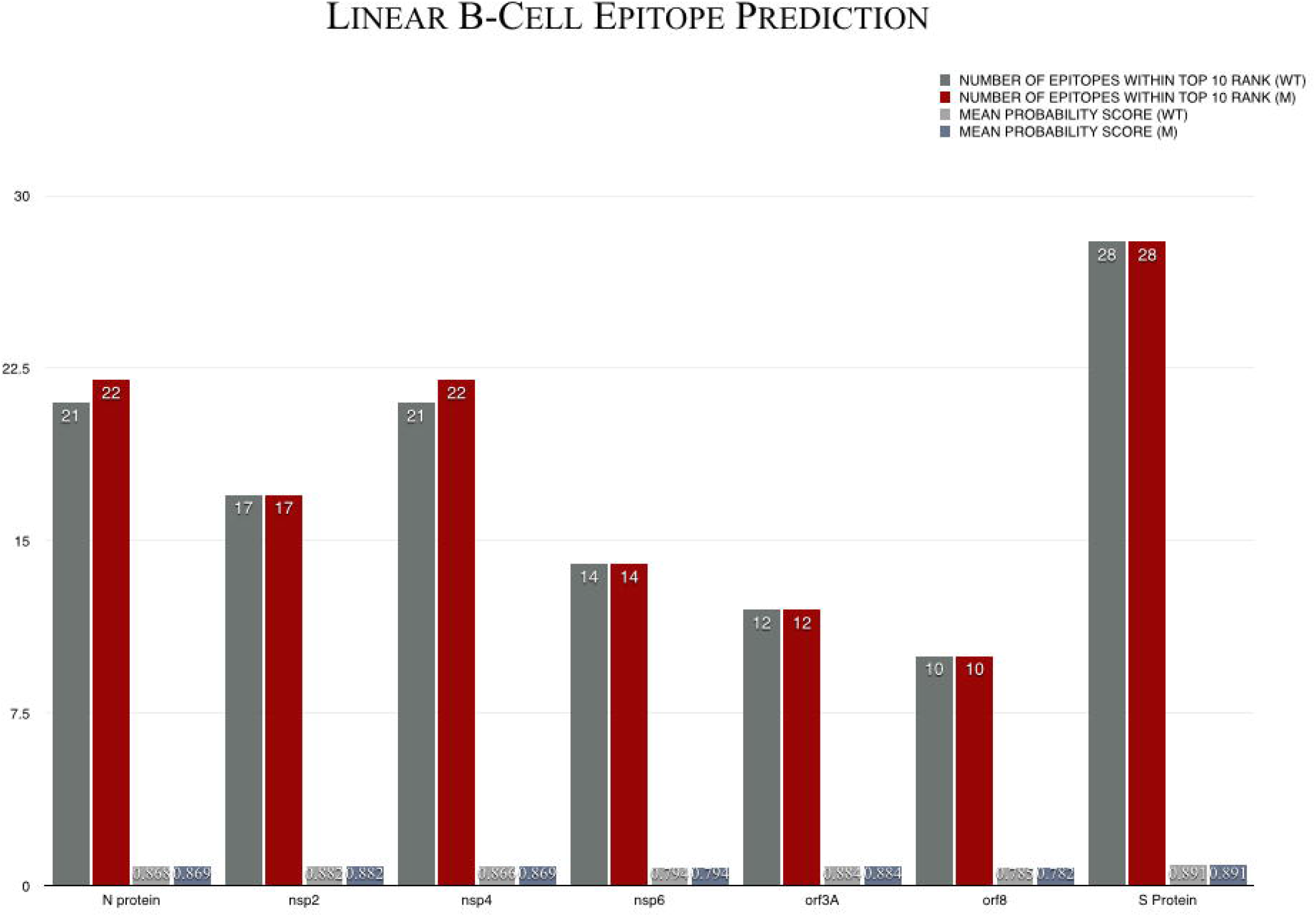
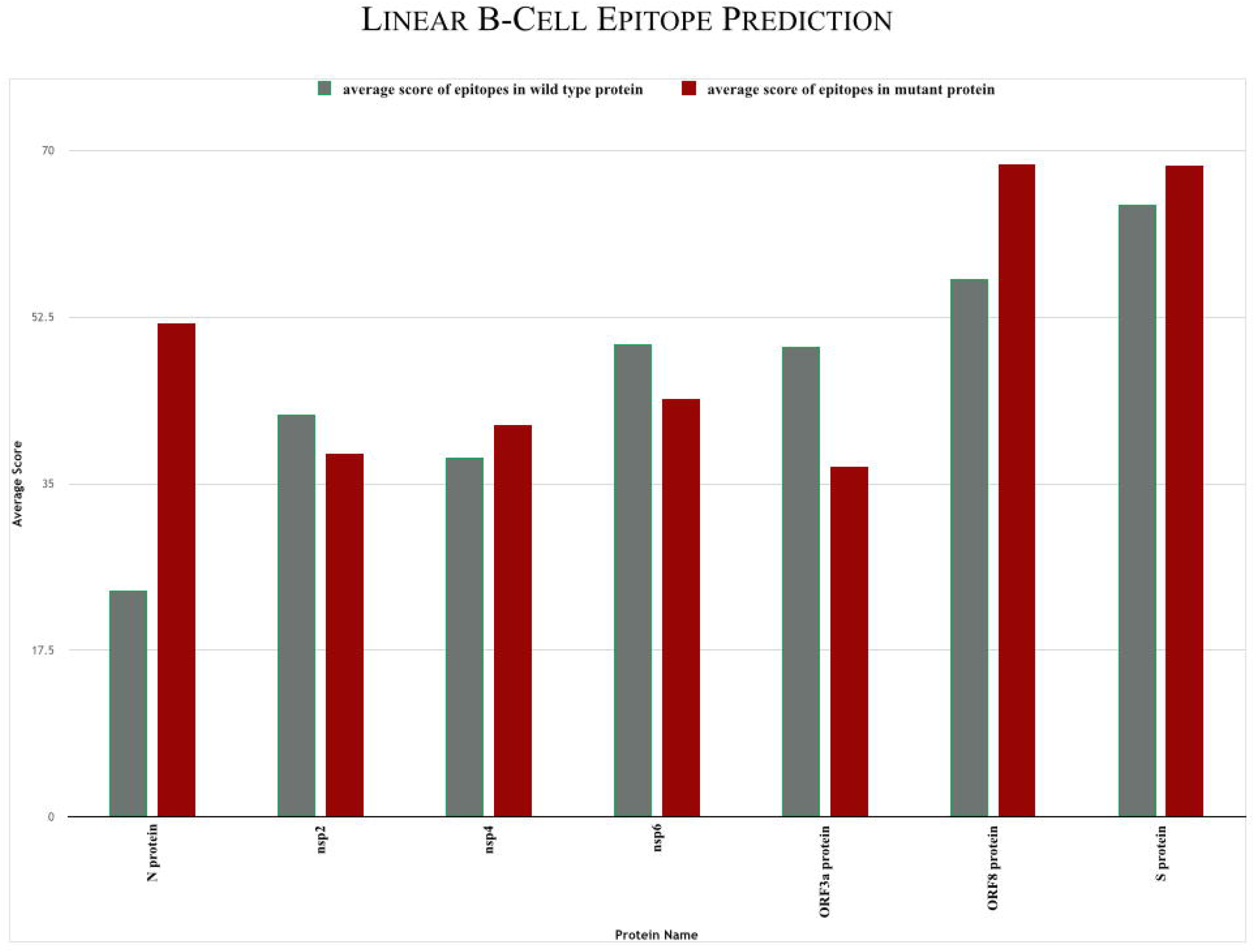
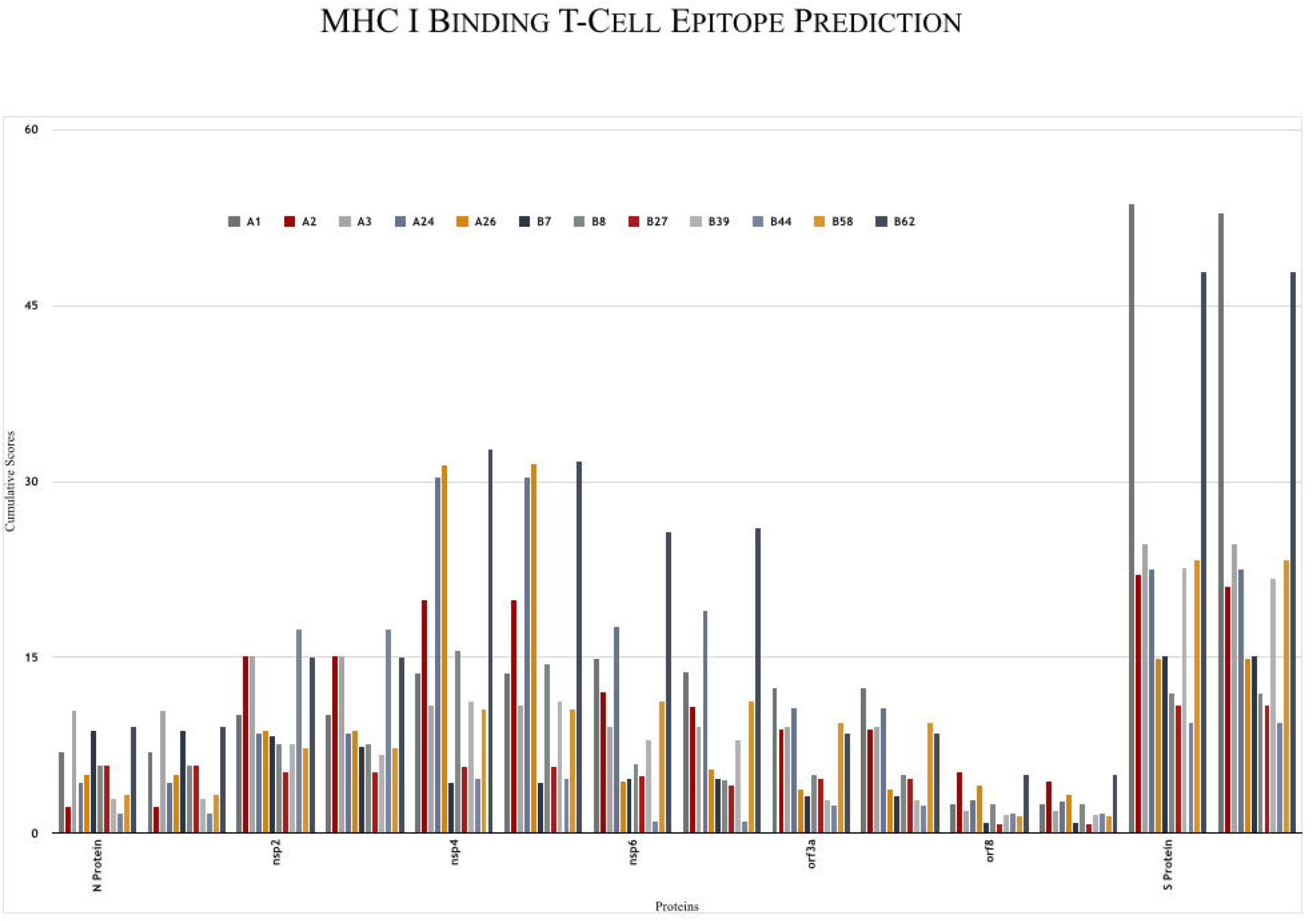
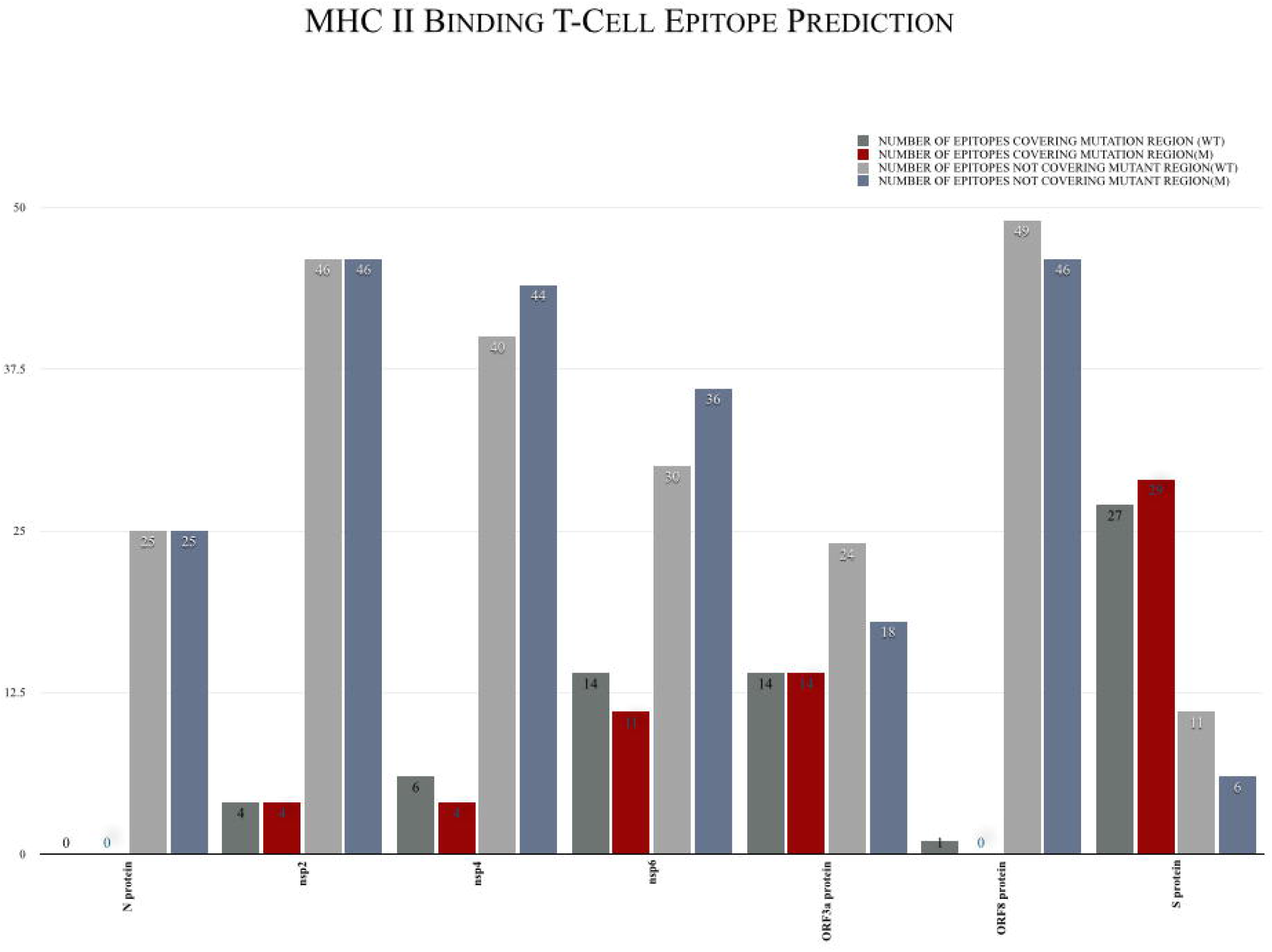
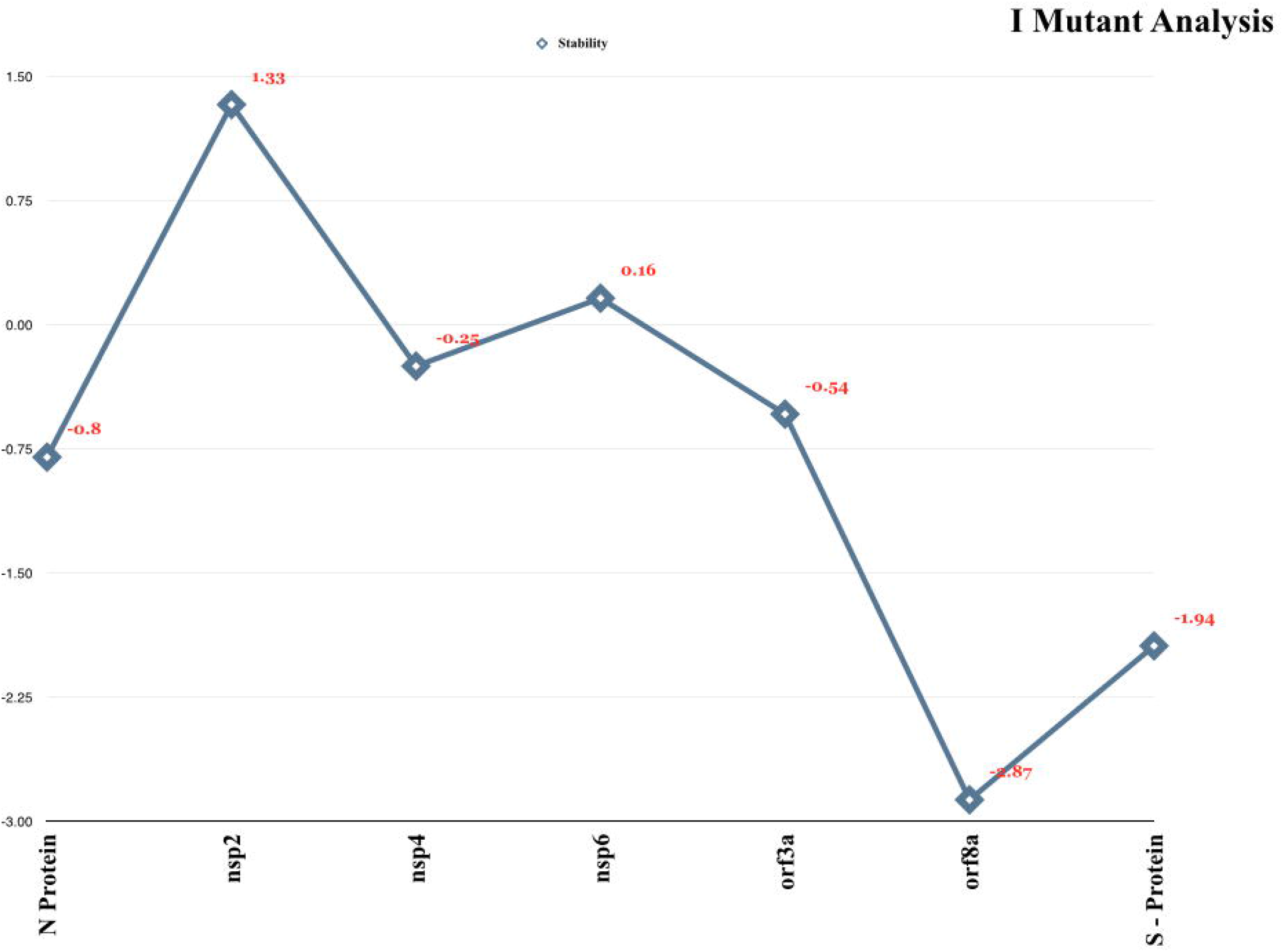
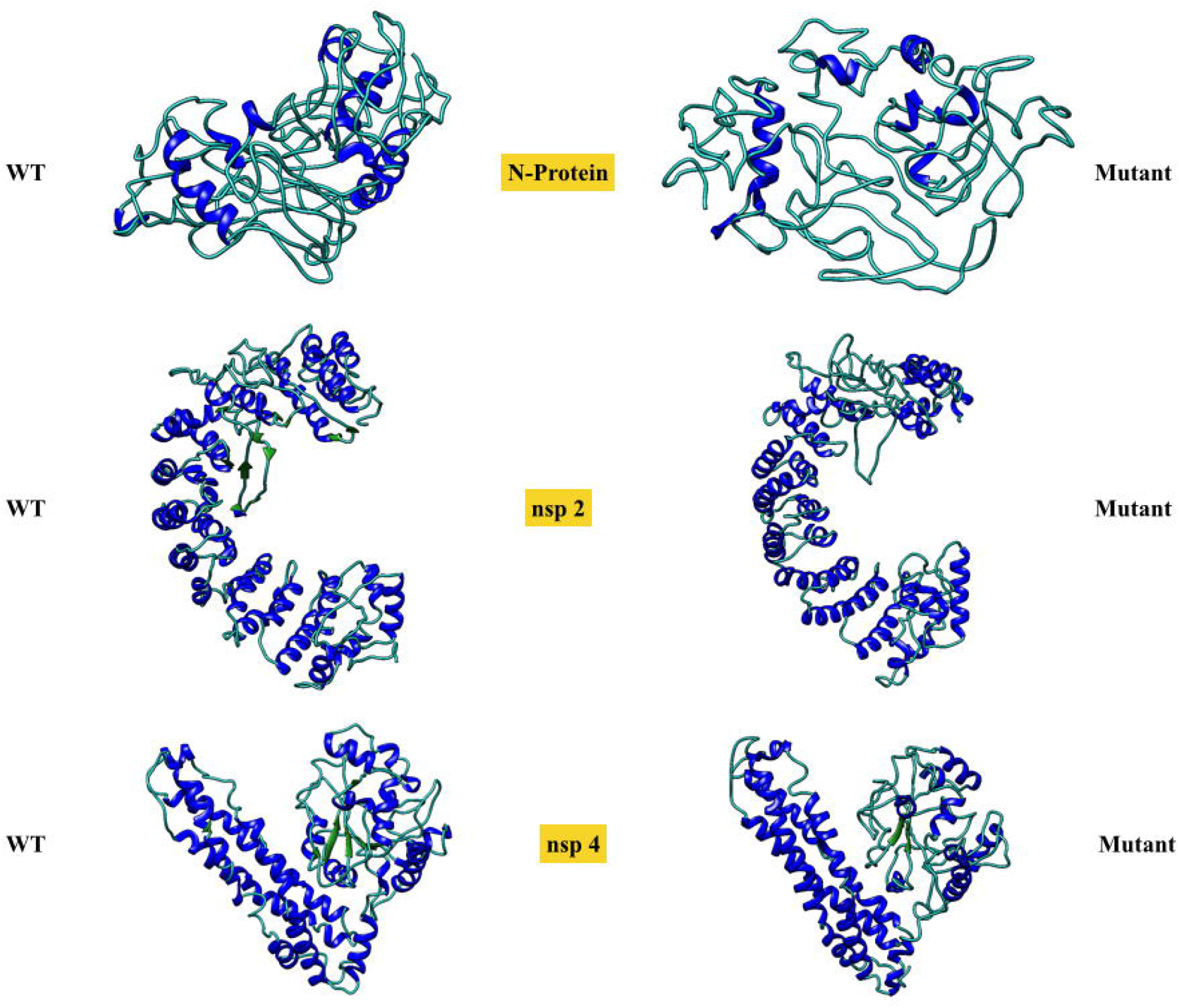
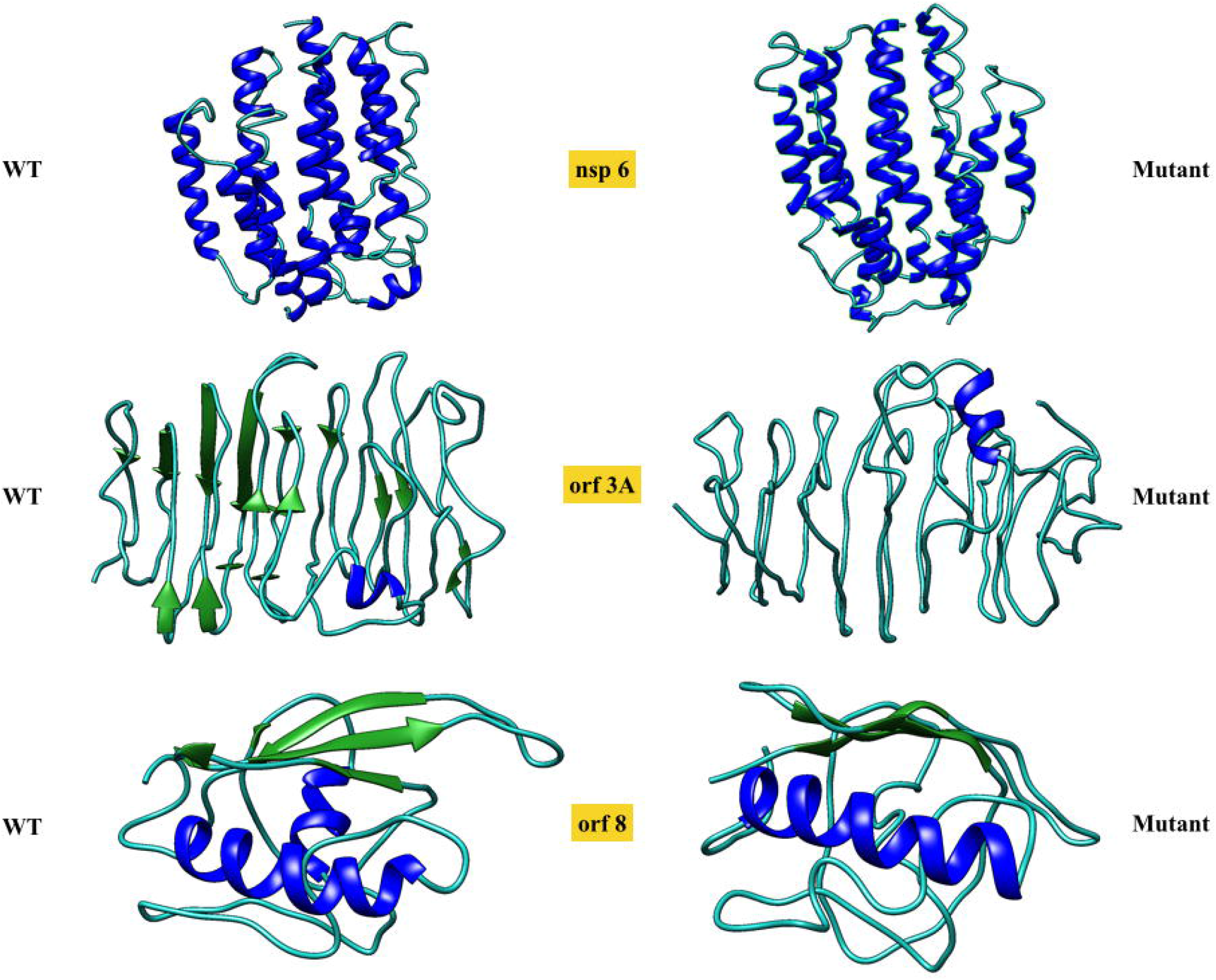
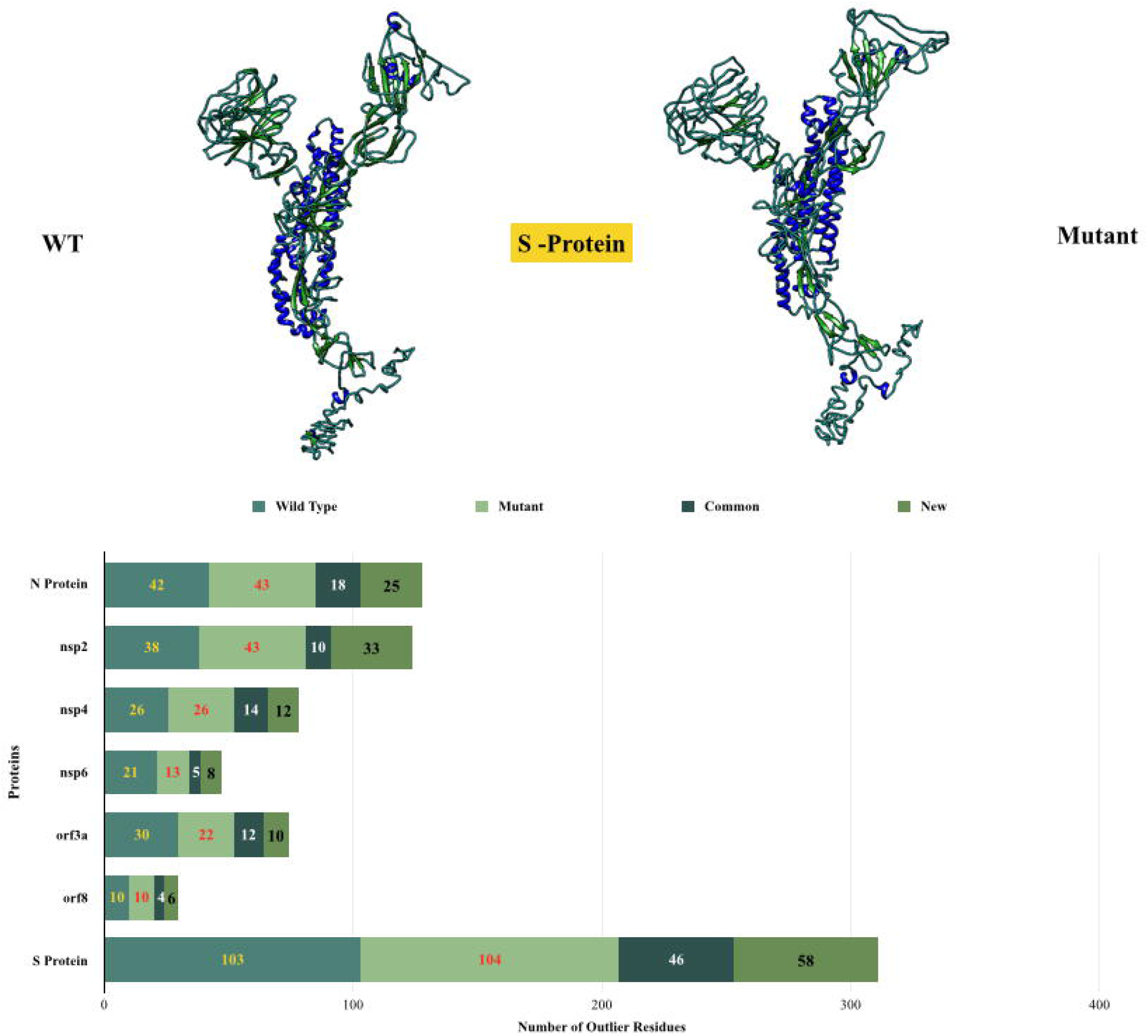
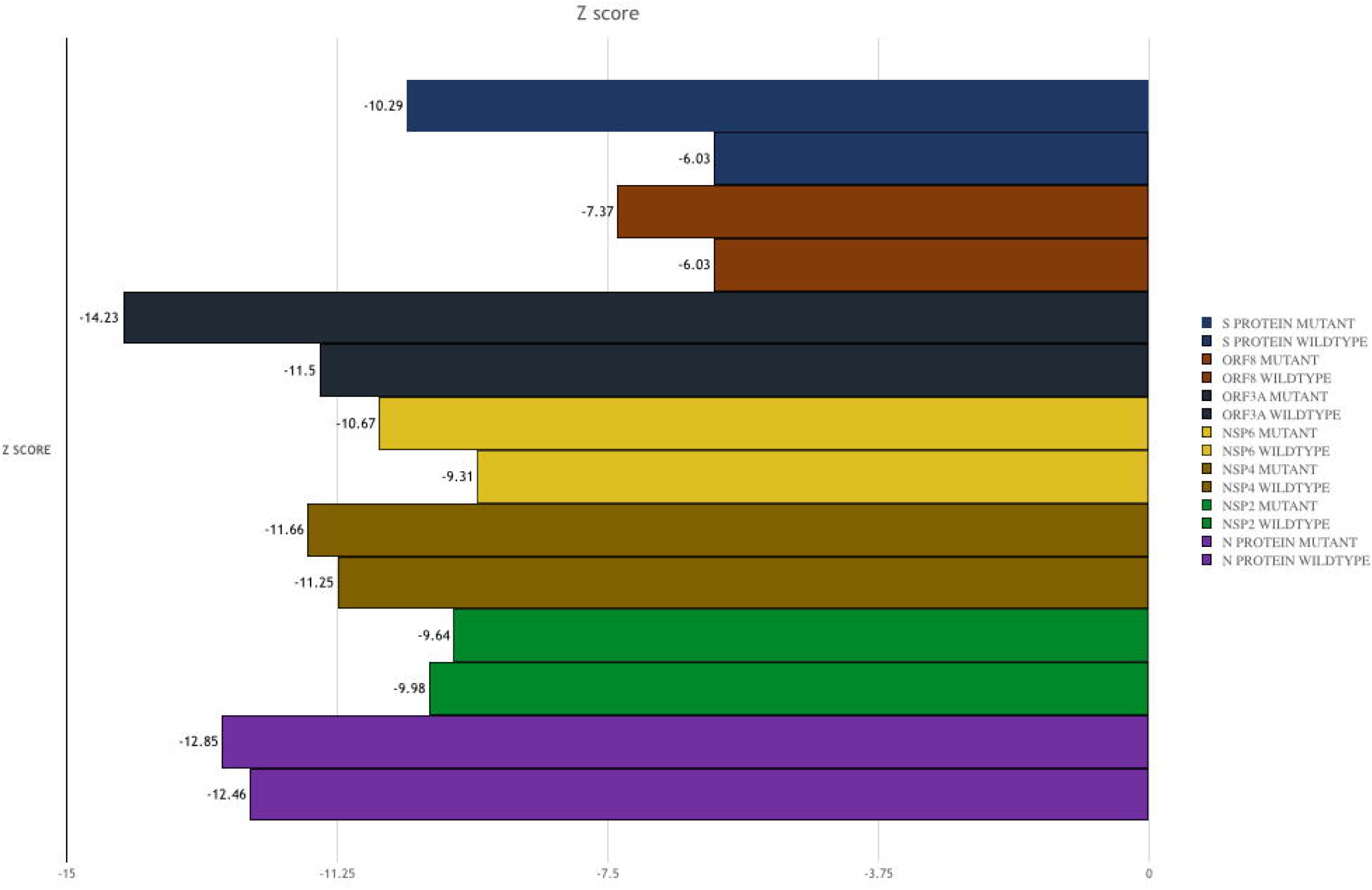
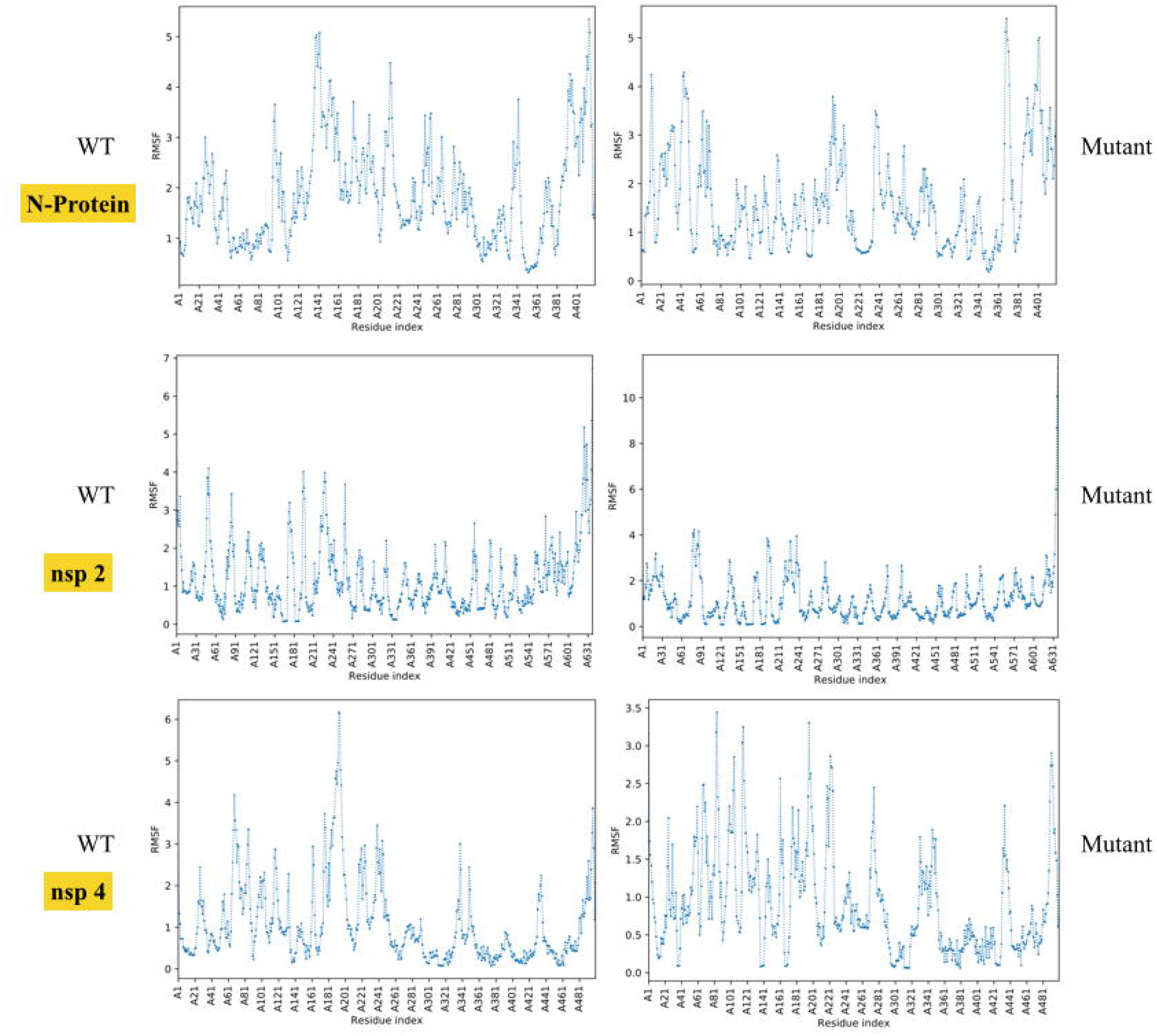
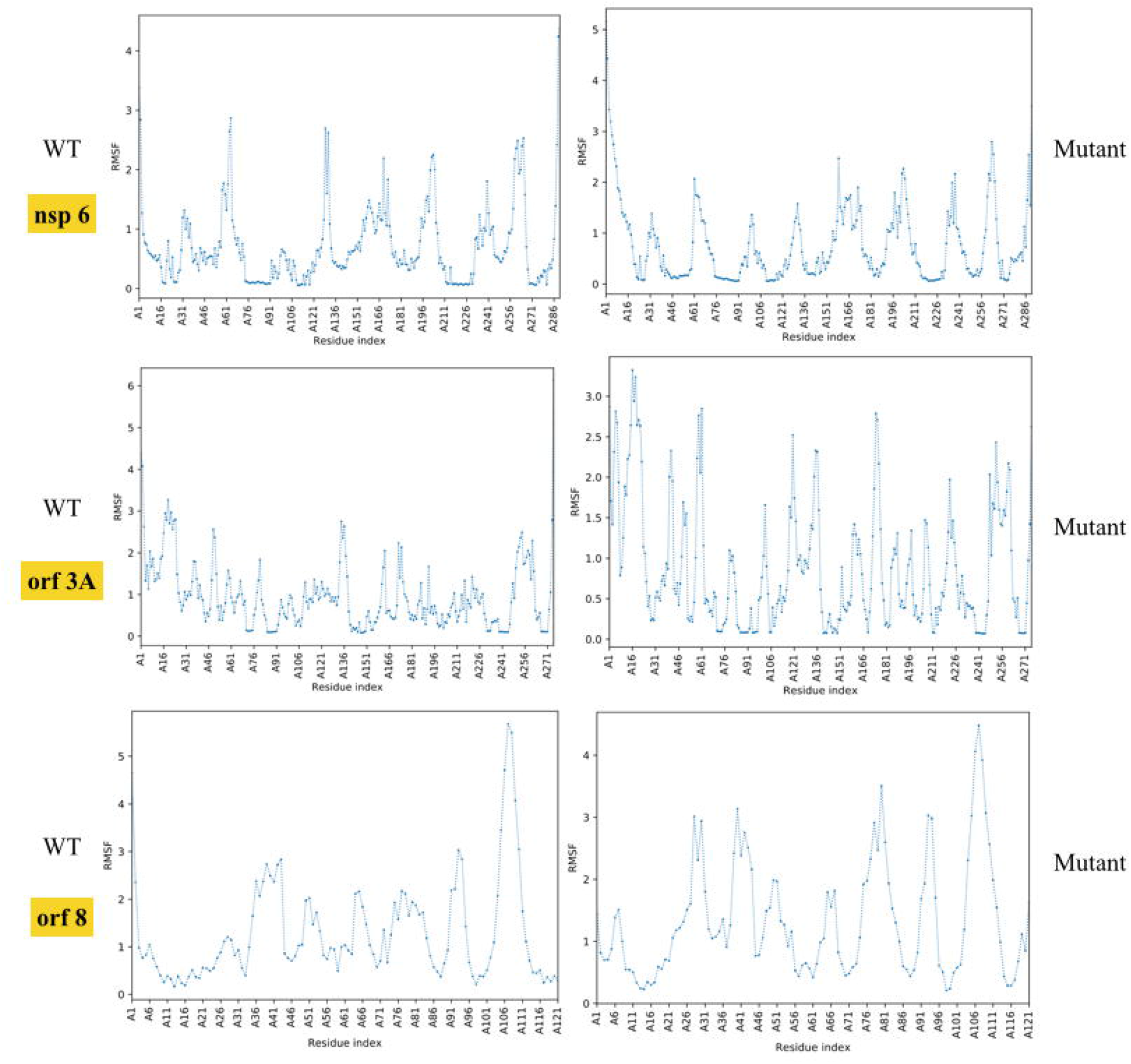
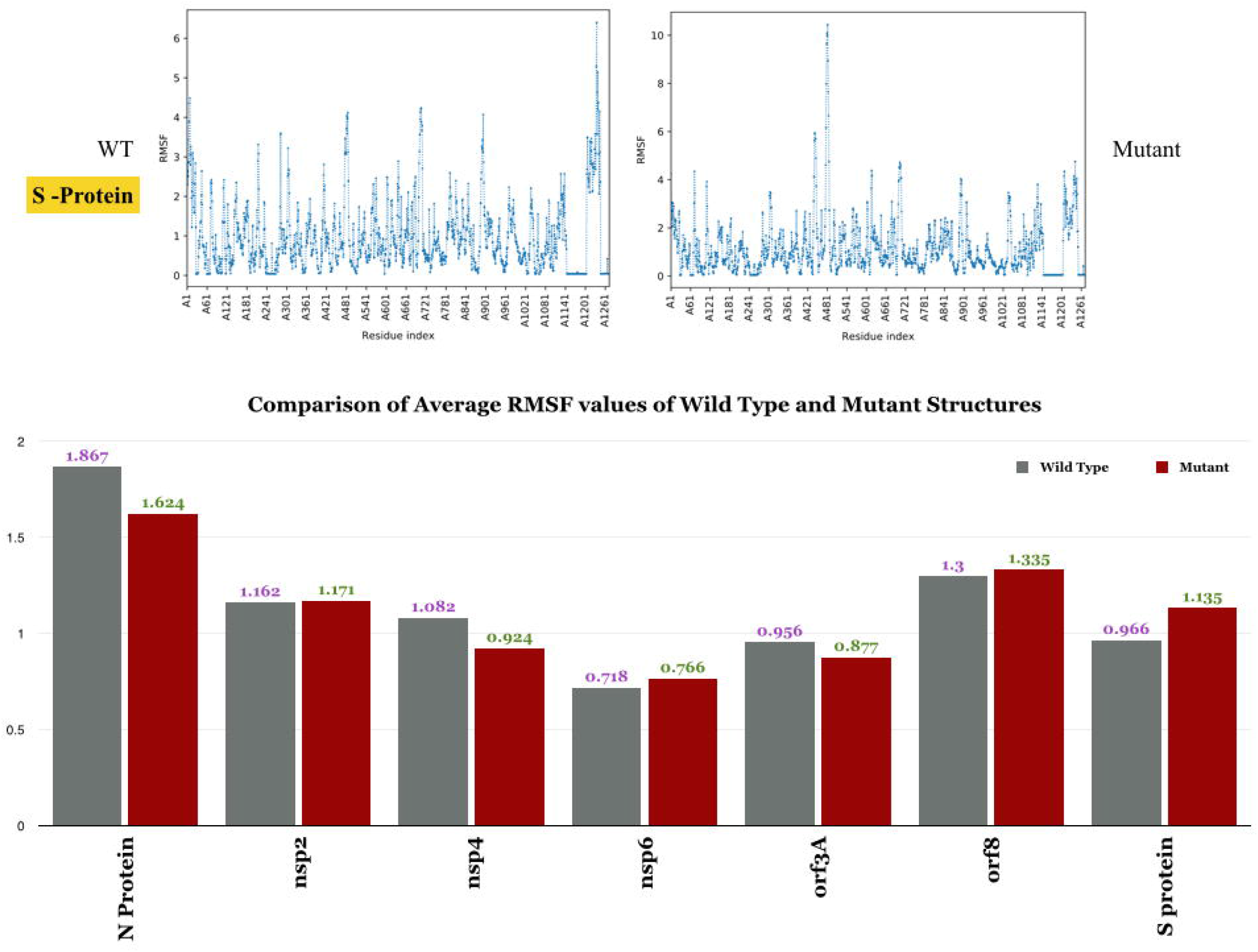
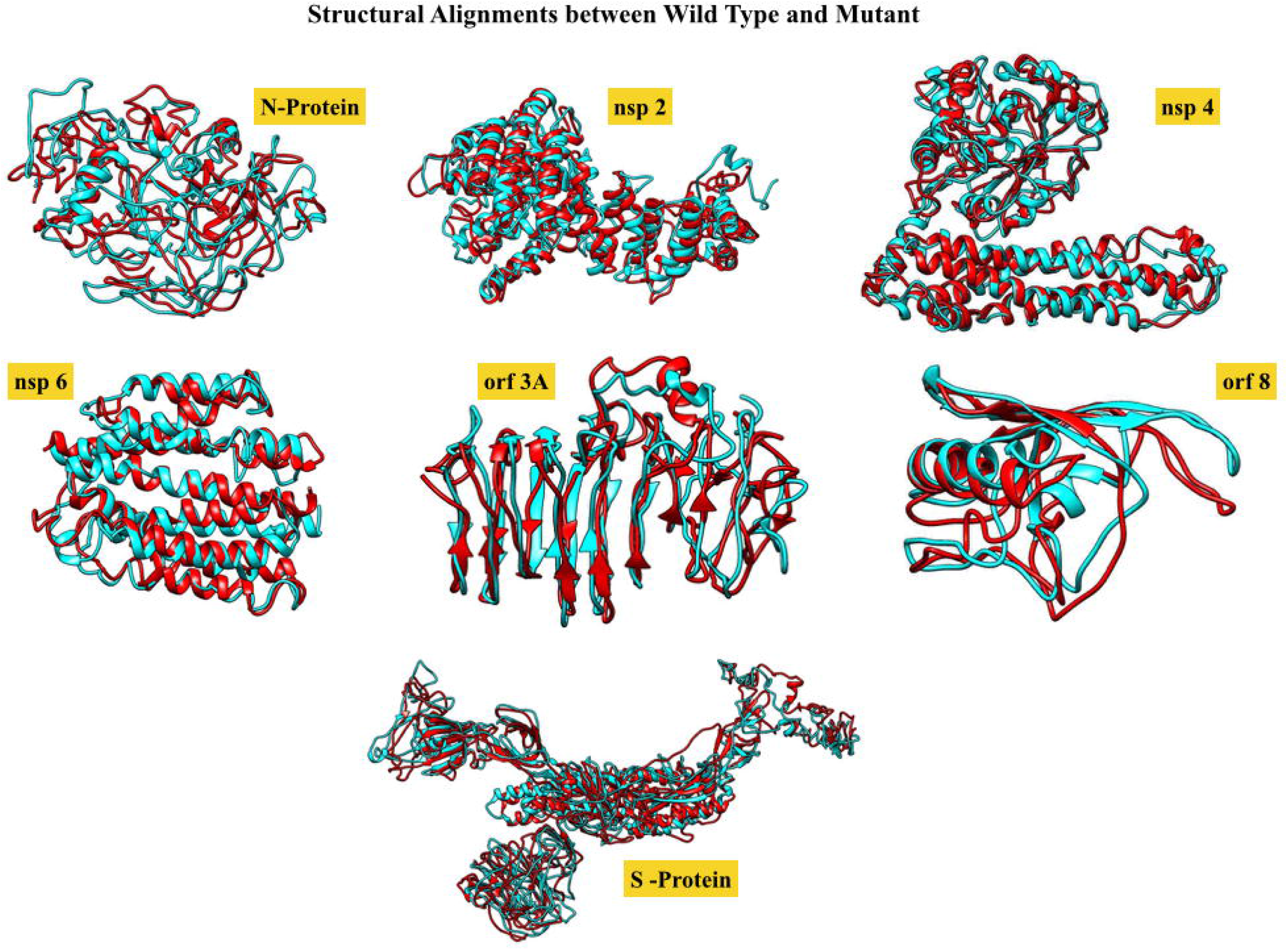
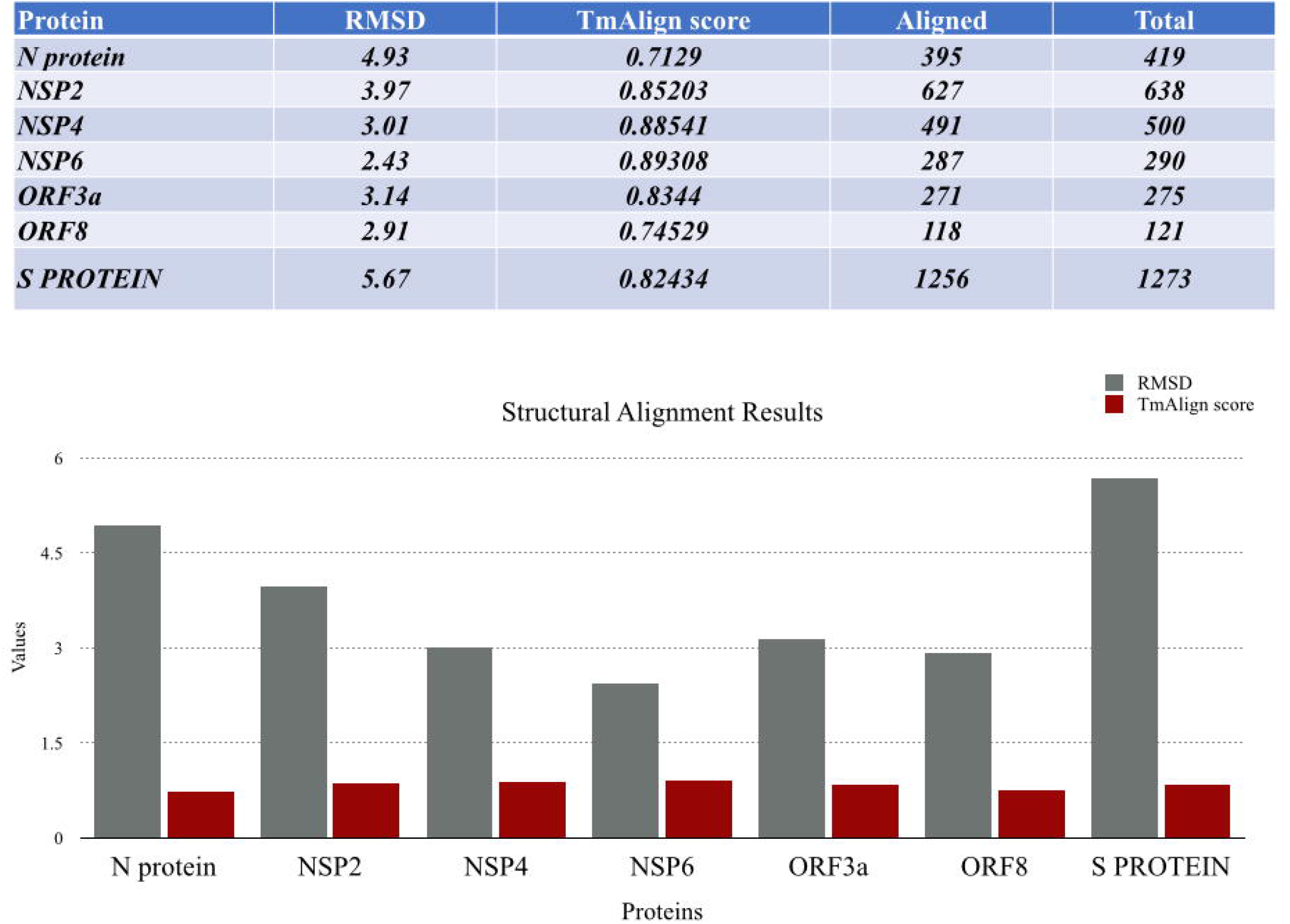
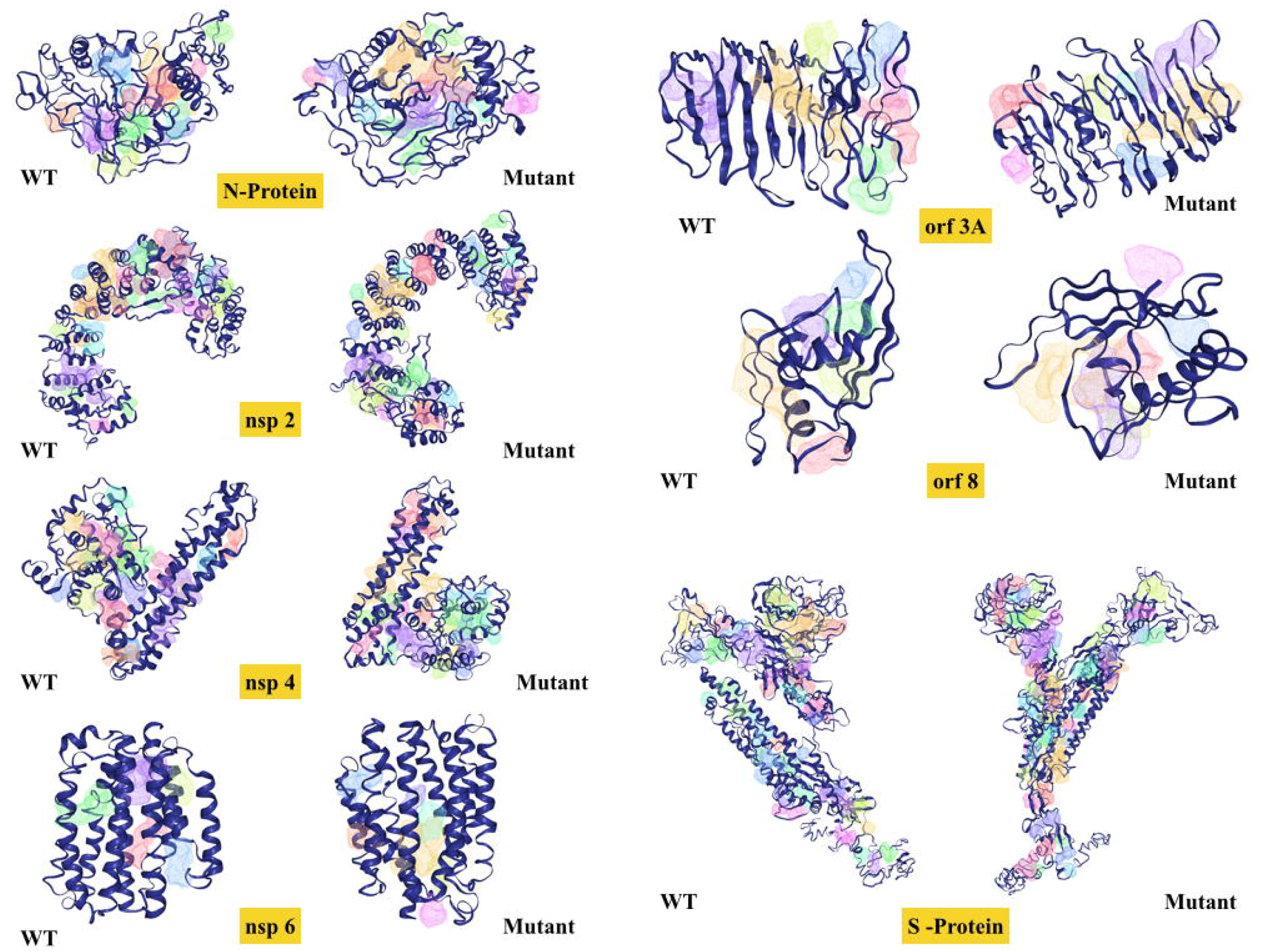
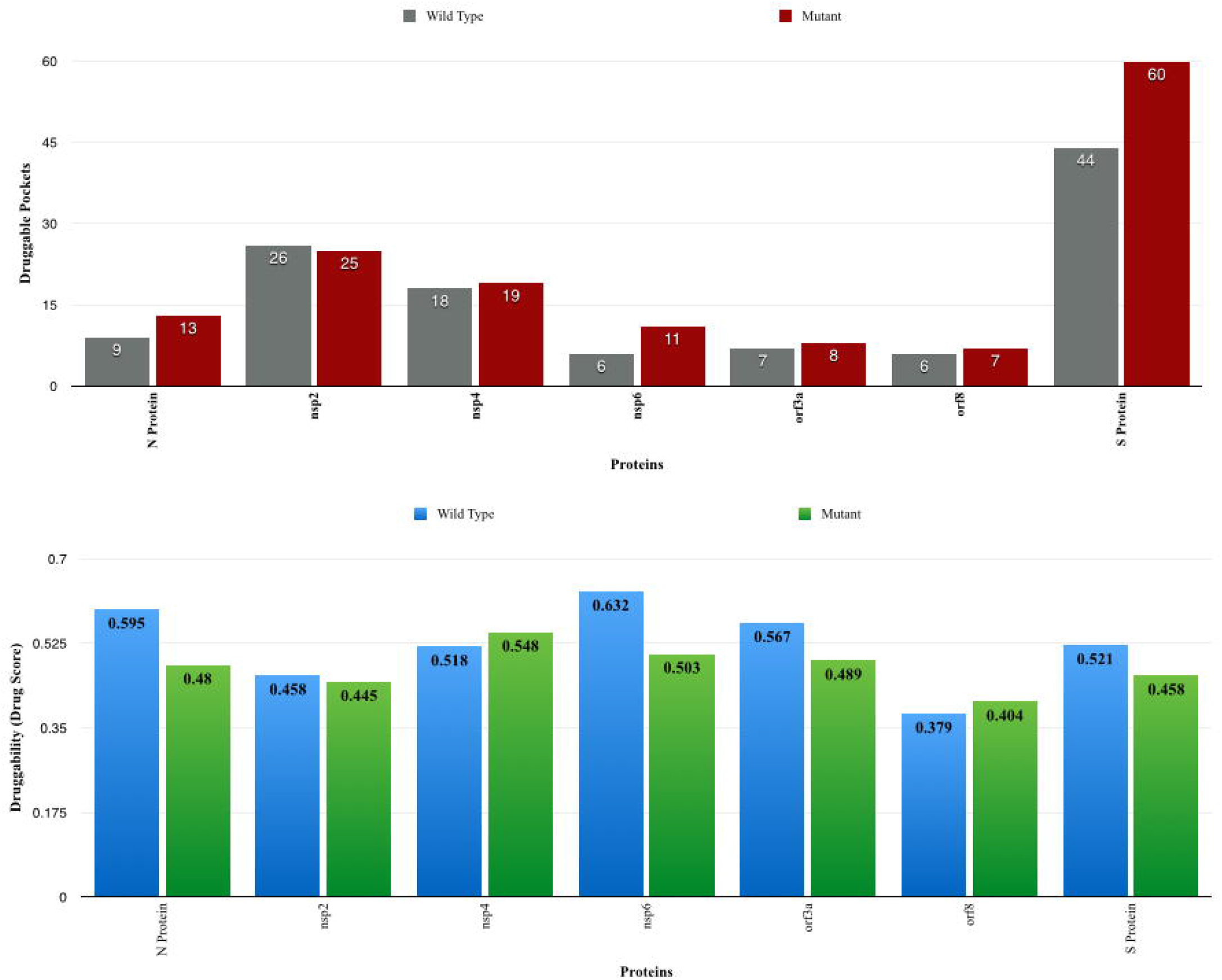
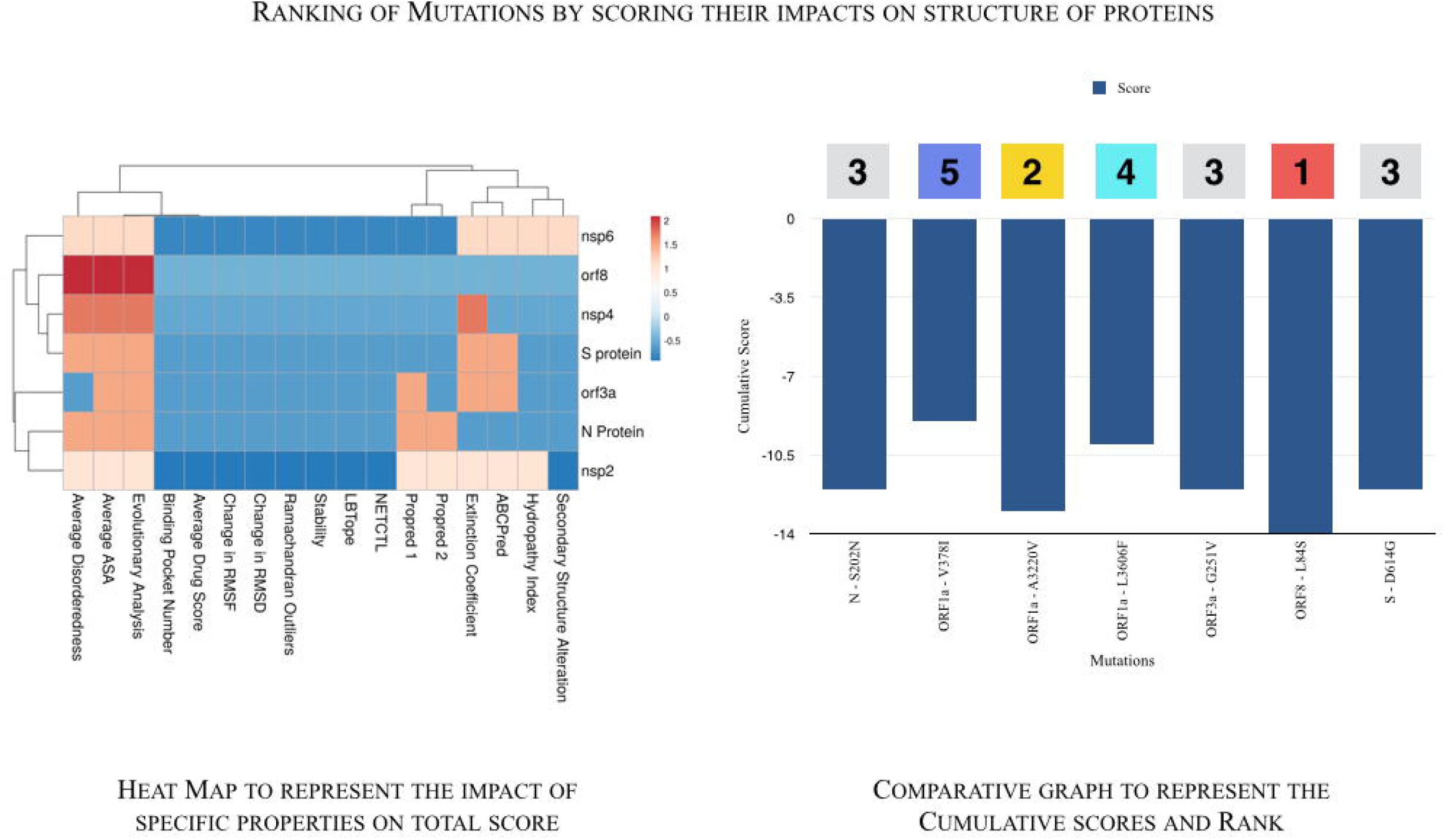

