## Supplementary figures and images for "Impact of clade specific mutations on structural fidelity of SARS-CoV-2 proteins"

### Ramachandran Plots Generated

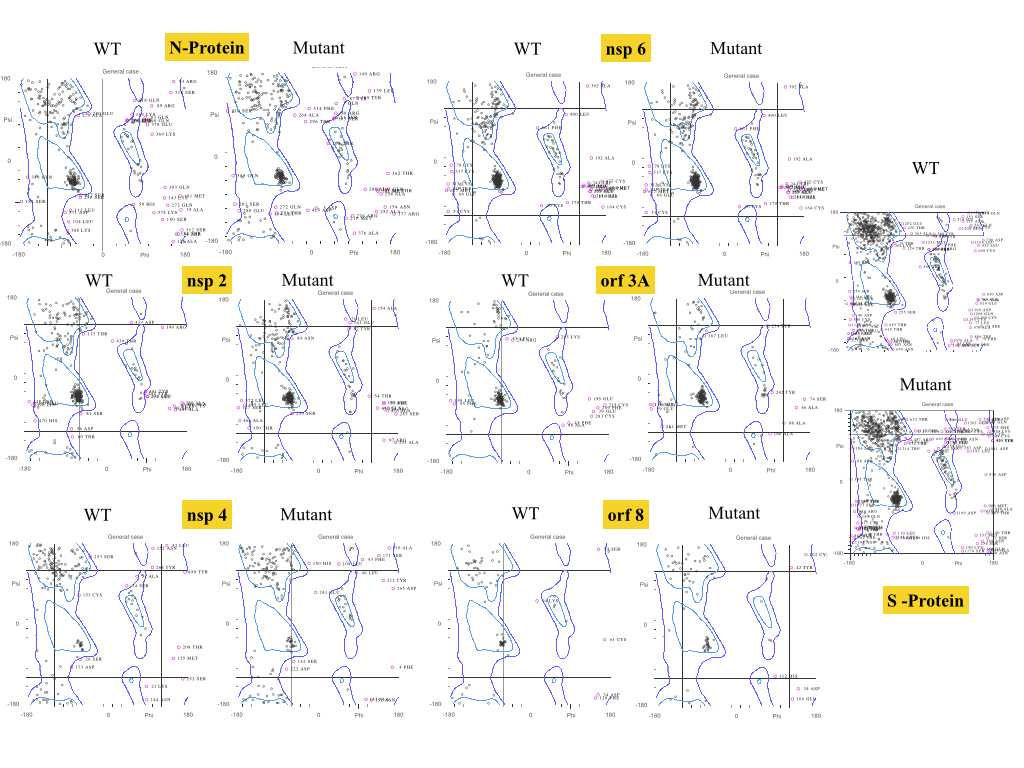
